# Identification of a novel HKU4-related coronavirus in single-cell datasets and clade viral host analysis

**DOI:** 10.1101/2023.06.18.545480

**Authors:** Adrian Jones, Steven E. Massey, Louis R. Nemzer, Daoyu Zhang, Yuri Deigin, Steven C. Quay

## Abstract

HKU4-related coronaviruses (CoVs) are merbecoviruses related to Middle Eastern Respiratory Syndrome coronavirus (MERS-CoV). In 2022 and 2023, two HKU4-related CoV strains were discovered in *Manis javanica* (Malayan pangolin) metagenomic datasets derived from organ samples: HKU4-P251T and MjHKU4r-CoV-1. Together with the *Tylonycteris robustula* bat CoV 162275, which was discovered in 2022, pangolin CoVs HKU4-P251T and MjHKU4r-CoV-1 form a novel phylogenetic clade distinct from all previously documented HKU4-related CoVs. In this study, we identified a novel HKU4-related CoV in a pangolin single-cell sequencing dataset generated by BGI-Shenzhen in Shenzhen, Guangdong, China in 2020. The CoV phylogenetically belongs to the same newly identified clade. The single cell datasets were reported as generated from organ samples of a single pangolin that died of natural causes. 98% of the HKU4-related CoV reads were found in only one of the seven single cell datasets — a large intestine cell dataset, cells of which exhibit low expression of DPP4. Bacterial contamination was found to be moderately correlated with HKU4-related CoV presence. We further identified with high confidence that the RNA-Seq dataset supporting one of four near identical variants of MjHKU4r-CoV-1 is a *Sus scrofa* (wild pig) metagenomic dataset, with only a trace level of *Manis javanica* genomic content. The presence of HKU4-related CoV reads in the dataset are almost certainly laboratory research-related and not from a premortal pangolin or pig infection. Our findings raise concerns about the provenance of the novel HKU4-related CoV we identify here, MjHKU4r-CoV-1 and its four near-identical variants.

## Introduction

The SARS epidemic in 2003-2004, as well as sporadic MERS outbreaks since the first case in 2012 (Zaki et al., 2012), have led to a significant increase in worldwide research of these human pathogenic coronaviruses (CoVs). Extensive sampling of bats for SARS-related and MERS-related CoVs, along with laboratory research on these viruses, ensued (Hu et al., 2017; Hu et al., 2018; Lattine et al., 2020). The source of the initial human SARS-CoV infection is thought to be masked palm civets in Guangdong markets (Hu and Shi, 2008), while MERS-CoV was transmitted to humans from camels on the Arabian peninsula (Alagaili et al., 2014). Bats were identified as the natural reservoir hosts of both SARS-related CoVs (Wang et al., 2006) and MERS-related CoVs (Wang et al., 2014). In 2019, SARS-CoV-2, a Sarbecovirus with an 82% identity to SARS-CoV, emerged in Wuhan. The origin of the virus has been the subject of an ongoing scientific debate and two main hypotheses have arisen. One hypothesis is that SARS-CoV-2 spilled over from animals at a seafood market in Wuhan (Zhou et al., 2020; Worobey et al., 2022), a second hypothesis is that the virus was accidentally released from a laboratory in Wuhan conducting SARS-related CoV research (Segreto and Deigin, 2020; Harrison and Sachs, 2022; Bahry, 2023).

HKU4-related CoVs, together with HKU5-related CoVs and MERS-related CoVs, belong to the Merbecovirus subgenus (previously called lineage C) of the betacoronavirus genus of coronaviruses (Fig. 1). HKU4-CoV and MERS-CoV use the dipeptidyl peptidase 4 (DPP4) receptor for cell entry.

**Fig. 1.**
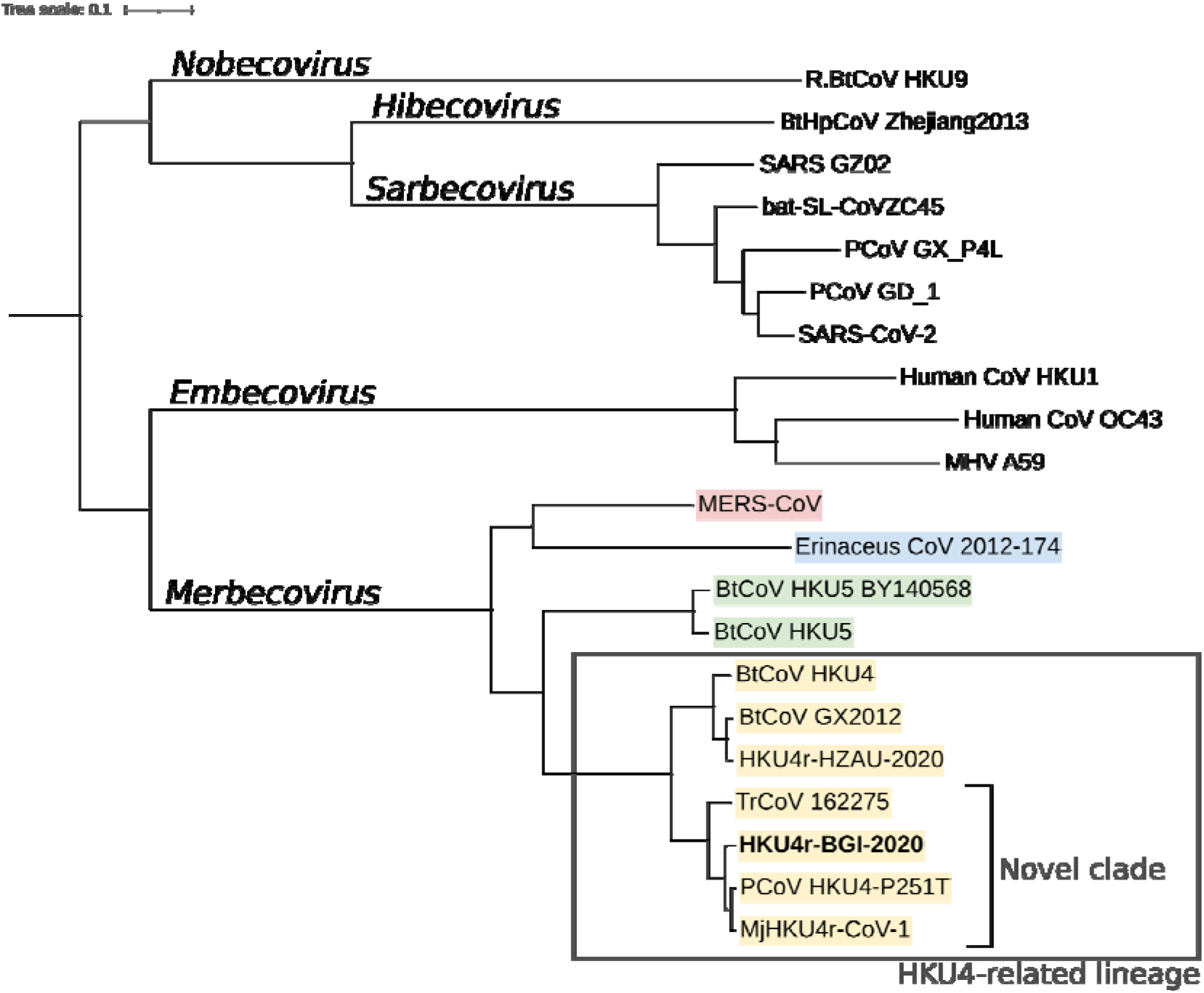
Betacoronavirus phylogenetic tree based on full genome sequences. The scale bar represents 0.1 substitutions per nucleotide position. The novel HKU4-related CoV identified here is shown in bold. Merbecoviruses are coloured by lineage. The tree is rooted on the midpoint and plotted using iTOL (Letunic and Bork, 2021).

In late 2019, a SARS-CoV-2 related partial genome was identified in pangolin tissue sequencing datasets (Liu et al., 2019). After the outbreak of COVID-19 in early 2020, two SARS-CoV-2-related strains, the Guangdong (GD) and Guangxi (GX) strains were identified in pangolin tissue samples. The GD pangolin CoV (PCoV) was identified by several authors (Zhang T. et al., 2020; Liu et al., 2020; Xiao et al., 2020; and Lam et al., 2020) who used next generation sequencing (NGS) datasets from Liu et al. (2019), as well as additional sequencing from the same batch of pangolins (Chan and Zhan, 2020). The GD pangolin coronavirus strain is particularly notable, as the amino acid sequence of the receptor binding domain (RBD) has a 97% identity to SARS-CoV-2 (Xiao et al., 2020; Lam et al., 2020; Liu et al., 2020; Zhang, T. et al., 2020). The Liu et al. (2019) sequencing datasets however, have been shown to be problematic. Hassanin (2020) identified human, mouse, and tiger genomic content contaminating the datasets. In addition, Jones et al. (2022c, 2022a) further noted the low GD PCoV read counts (0.16 to 30 reads per million reads), a notable correlation between bacterial content and GD PCoV read count (Spearman’s rho of 0.58), and human and mouse genomic sequences contaminating the datasets sequenced by Liu (2019). Noting a 10× higher binding affinity of GD PCoV for human ACE2 than to pangolin ACE2 (Wrobel et al., 2021), Jones et al. (2022c, 2022a) raise the possibility that the SARS-CoV-2-related CoV in Liu et al. (2019) may stem from laboratory contamination.

GX pangolin coronaviruses were first identified by Lam et al. (2020) in frozen *Manis javanica* (Malayan pangolin) tissue samples collected in 2017/2018 by the Guangxi Zhuang Autonomous Region Terrestrial Wildlife Medical-aid and Monitoring Epidemic Diseases Research Center. Peng et al. (2021) identified a partial GX PCoV sequence, MP20 in frozen *Manis pentadactyla* (Chinese pangolin) tissue sampled in Yunnan province in 2017. The only other documented recovery of GX PCoV sequences in pangolins was by Nga et al. (2022), who identified partial RdRp and NSP14 sequences related to GX PCoVs in *M. javanica* tissue samples collected in Vietnam. A GX PCoV related partial genome, GX_ZC45r-CoV was recovered by Jones et al. (2022b) from sequence read archive (SRA) datasets from seven animal species collected across China and sequenced by He et al. (2022). The partial genome exhibits an ancestral phylogenetic relationship to GX PCoVs in the NSP4, NSP10, and partial RdRp regions, but also exhibits close grouping with Zhoushan bat CoV bat-SL-CoVZC45 (Hu et al., 2018) for the full RdRp gene.

Interestingly, both the GD and GX strains appear to be rare, with no infection in the wild identified in 334 samples from *M. javanica* pangolins in Malaysia (Lee et al., 2020), nor in 33 *M. javanica* and *M. pentadactyla* species sampled in Zhejiang between 2017 and 2020 (He et al., 2022). This is true among 161 pangolins trafficked in the 2018 to 2019 period from Southeast Asia into China (Shi et al., 2022), in the 93 pangolin samples sampled between 1990 to 2017 from pangolins seized in Guangzhou, and Yunnan provinces in China, Malaysia, Myanmar and Taiwan (Hu et al., 2020; Zhang 2020), in 17 pangolins collected from undisclosed locations in China between November 2019 and March 2020 (Deng et al., 2020), in seven *M. javanica* collected by Guangdong Provincial Wildlife Rescue Centre from unknown locations or dates (Li et al., 2020), and in three pangolins sampled in Guangxi province by Wenzhou University in May through July 2020 (Jones et al., 2020c). GD, GX and HKU4-related CoVs detected in pangolin RNA-Seq data and PCR products are listed in Supp. Info. 1.

In August 2022, Shi et al. (2022) reported results from the sequencing of tissues from 161 trafficked *M. javanica* pangolins in China. Shi et al. identified a novel HKU4-related CoV, PCoV HKU4-P251T, in one pangolin. This was the first reported merbecovirus identified in pangolin tissues. In February 2023, Chen el al. (2023) identified a similar HKU4-related CoV (MjHKU4r-CoV-1 and variants) in four out of 86 pangolins seized by Guangxi customs, and smuggled from “Southeast Asian countries” (Chen et al., 2023). Anal swab samples for the four pangolins with a HKU4-related CoV were all collected on 2019-11-08. In April 2023, Cui et al. (2023) documented the identification of HKU4 isolate GX/HKU4-GX/2020, nearly identical to MjHKU4r-CoV-1 in a *M. javanica* pangolin sampled in Guangxi province, China in 2020.

Prior to the published discovery of HKU4-related CoVs in pangolin RNA-Seq datasets, in early 2020, Chen et al. (2020) undertook single-cell screening of SARS-CoV-2 target cell lines of 11 species, including *Felis catus* (domestic cat), *Sus scrofa domesticus* (pig), and a single *M. javanica* pangolin (Supp. Info. 2.1). The research was initially disclosed as a preprint in June 2020, and later published in Chen et al. (2021). The single wild pangolin whose tissues were used for analysis died of natural causes, and was sourced from the Guangdong Provincial Wildlife Rescue Center, while other animals were sourced from pet and agricultural markets. Single-nuclei libraries for multiple tissues were generated, and the relative expression patterns of 114 cell receptors of 144 viruses, including the ACE2 receptor for SARS-CoV-2 and DPP4 for MERS, were quantified. The libraries were sequenced on the BGISEQ-500 platform and registered on NCBI on 2021-07-18 as BioProject PRJNA747757, and published on NCBI between the 2021-07-22 and 2021-09-02 by BGI-Shenzhen. Surprisingly, pangolins were assessed to be poorly susceptible to SARS-related CoVs, with only pangolin kidney proximal tubule cells inferred as susceptible, due to receptor expression levels. However a variety of pangolin cells were inferred as susceptible to MERS-related CoVs including lung, kidney, and spleen cells.

In the present study, we identified a novel HKU4-related CoV in seven pangolin single cell sequencing RNA-Seq datasets generated by Chen et al. (2021). We identified that the HKU4-related CoV groups phylogenetically with three recently published HKU4-related CoVs to form a novel clade. We analyzed genome and amino acid similarity to related CoVs in the novel clade. We further reviewed mammalian genomic, viral and bacterial contamination of all datasets generated by Chen et al. (2021). We reviewed three scenarios to explain the presence of the novel HKU4-related CoV in the single cell RNA-Seq datasets sequenced by Chen et al. (2021) and conclude that its presence is related to laboratory research rather than naturally infected pangolin organs.

Finally, we undertook metagenomic analysis of selected datasets published by Cui et al. (2023) containing an RNA-Seq dataset for HKU4 isolate GX/HKU4-GX/2020, a near identical variant of MjHKU4r-CoV-1. The HKU4-GX dataset was reported to have been sampled from a *M. javanica* specimen. However we identified this sample as a *S. scrofa* sample with only trace levels of *M. javanica* genomic content. We conclude the presence of the CoV in the dataset could not have been sourced from the trace level of *Manis* genomic content and is, by inference, laboratory research-related.

## Results

### Novel HKU4-related CoV

A HKU4-related coronavirus (CoV) was initially identified in three *Manis javanica* cell sequencing datasets in BioProject PRJNA747757 obtained by Chen et al. (2021), using fastv and NCBI STAT. Pangolin large intestine dataset PGN_LI was found to have the highest coverage of the *Tylonycteris* bat coronavirus HKU4 genome. We aligned the PGN_LI dataset to the *Tylonycteris bat* coronavirus HKU4 genome, and then analyzed a consensus sequence using blastn. The highest homology of the consensus genome at the time of analysis was to PCoV HKU4-P251T, a genome published on NCBI on 2022-02-06. We then aligned each sequence read archive (SRA) dataset in BioProject PRJNA747757 to PCoV HKU4-P251T and identified the novel HKU4-related CoV in two additional pangolin cell datasets, lung dataset PGN_LG and esophagus dataset PGN_ES. We pooled the three SRA datasets, and performed *de novo* assembly of the reads using MEGAHIT. We then aligned both the pooled reads from PGN_LI, PGN_LG and PGN_ES to PCoV HKU4-P251T using minimap2. A preliminary consensus genome was then generated from aligned reads and *de novo* assembled contig sequences.

To extract maximum read coverage, we infilled gaps in coverage of the novel HKU4-related CoV preliminary consensus genome using the PCoV HKU4-P251T genome and aligned all SRA datasets in BioProject PRJNA747757 to this infilled genome. Seven pangolin cell datasets in total were identified as containing HKU4-related CoV reads. A final consensus novel HKU4-related CoV genome was then generated as described in Methods.

We then analyzed the resulting consensus sequence using blastn (Table 1). Genome coverage is incomplete at 85%, with 97.34% and 97.23% identities to pangolin CoVs MjHKU4r-CoV-1 and PCoV HKU4-P251T, respectively. This is of note, as MjHKU4r-CoV-1 and PCoV HKU4-P251T have a 99.06% overall identity to each other. The closest bat CoV is *Tylonycteris robustula* coronavirus isolate 162275, which was sampled in Yunnan in 2016, with a 92.96% identity. We refer to the novel HKU4-related CoV identified here as HKU4r-BGI-2020, to reflect its taxonomic affiliation, location, and year of sequencing.

**Table 1.** Top 10 results of blastn analysis of HKU4r-BGI-2020 against the nt database and the MjHKU4r-CoV-1 genome (Chen et al., 2023). See Supp. Info. 3 for accession numbers.

| Description | Max Score | Total Score | Query Cover | Per. ident |
| --- | --- | --- | --- | --- |
| MjHKU4r-CoV-1 | 4721 | 43485 | 85% | 97.34 |
| PCoV HKU4-P251T | 4704 | 43501 | 85% | 97.23 |
| TrCoV isolate 162275 | 4039 | 37966 | 84% | 92.86 |
| HKU4-related isolate GZ131656 | 2874 | 25049 | 72% | 85.29 |
| HKU4r-HZAU-2020 | 2852 | 26520 | 76% | 85.13 |
| BtTp-BetaCoV/GX2012 | 2841 | 26496 | 76% | 85.09 |
| BtCoV HKU4-3 | 2824 | 26301 | 75% | 84.93 |
| BtCoV HKU4-2 | 2824 | 26609 | 76% | 84.93 |
| T.BtCoV HKU4 | 2824 | 26301 | 75% | 84.93 |
| BtCoV HKU4-4 | 2819 | 26583 | 76% | 84.9 |
| T.BtCoV HKU4 isolate SM3A | 2813 | 26436 | 76% | 84.86 |

We note that MjHKU4r-CoV-1 (Chen et al., 2023) differs by only two nucleotides (3025/3027 nt) compared with HKU4 isolate GX/HKU4-GX/2020 (Cui et al., 2023). Since these two variants, as well as variants MjHKU4r-CoV-2-4, are nearly identical, we only included MjHKU4r-CoV-1 in further analyses.

98% of HKU4r-BGI-2020 aligned reads were found in the pangolin large intestine cell sample PGN_LI, with only a trace level of the reads aligning to the novel HKU4-related CoV in pangolin lung sample PGN_LG (Table 2).

**Table 2.** All pangolin cell sample datasets in BioProject PRJNA747757 were aligned to the HKU4r-BGI-2020 sequence, and alignment statistics tabulated including coverage percentages at one or more, 10× and 30× read depths. See Supp. Info. 2.3 for MA minimum cutoff counts.

| SRA | Sample | Organ | HKU4r Counts | Counts/M spots | Coverage% | Cov. 10x % | Cov. 30x % |
| --- | --- | --- | --- | --- | --- | --- | --- |
| SRR15199667 | PGN_LI | Large intestine | 51558 | 321.753 | 83.85 | 58.25 | 30.13 |
| SRR15199671 | PGN_ES | Esophagus | 838 | 8.877 | 18.19 | 1.7 | 1.22 |
| SRR15199672 | PGN_DM | Duodenum | 44 | 0.617 | 1.42 | 0.41 | 0 |
| SRR15199666 | PGN_LR | Liver | 7 | 0.115 | 1.22 | 0 | 0 |
| SRR15199668 | PGN_LG | Lung | 78 | 0.112 | 2.92 | 0.82 | 0.18 |
| SRR15199665 | PGN_SH | Stomach | 18 | 0.106 | 2.4 | 0 | 0 |
| SRR15199670 | PGN_HT | Heart | 1 | 0.010 | 0.23 | 0 | 0 |
| SRR15199663 | PGN_SN | Spleen | 0 | 0 | 0 | 0 | 0 |
| SRR15199669 | PGN_KY | Kidney | 0 | 0 | 0 | 0 | 0 |

All reads aligning to the HKU4r-BGI-2020 genome were pooled and alignments reviewed. A large number of reads are found aligning to the 3’ UTR region, with significant clusters of reads aligning to the 5’ UTR and NSP13 coding region (Fig. 2).

**Fig. 2.**
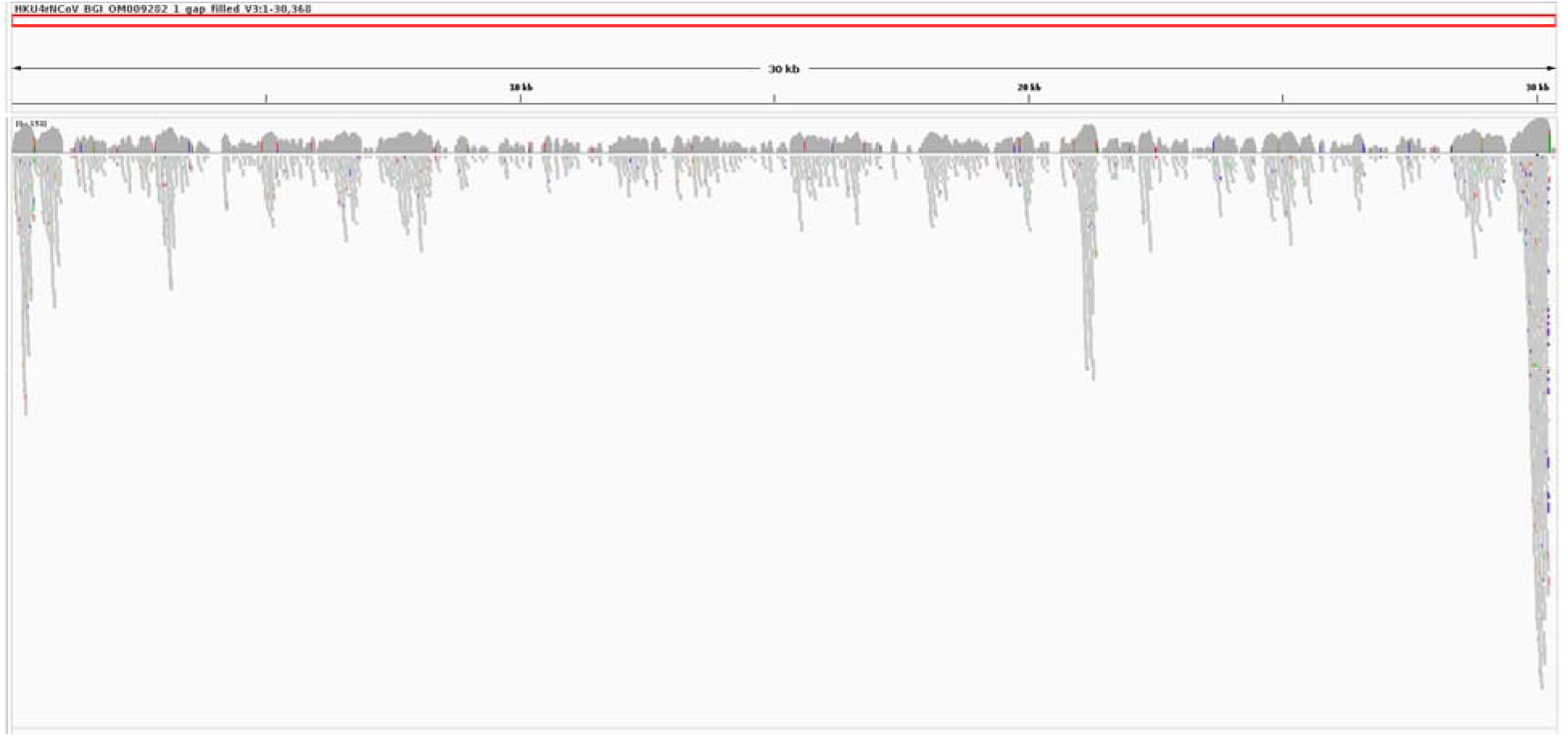
Reads mapping to HKU4r-BGI-2020 from 7 pangolin cell datasets. Aligned using minimap2. Reads of 28-nt length or shorter were excluded and only those meeting a minimum MAPQ score of 20 were plotted. Average read depth is 113 reads, with a median of 13 reads. Displayed using IGV.

### Similarity plot

Similarity plot analysis shows the novel HKU4r-BGI-2020 CoV to have the highest similarity to MjHKU4r-CoV-1 and PCoV HKU4-P251T across the genome. More divergence from these genomes occurs in two regions, in the Nsp13 region of ORF1ab and in the accessory protein genes ORF3a-d (encoding NS3a-d as per Woo et al. (2007)) (Fig. 3). Indeed, in the Nsp13 region of ORF1ab, HKU4r-BGI-2020 matches *Tylonycteris robustula* coronavirus isolate 162275 with a significantly higher identity than either MjHKU4r-CoV-1 or PCoV HKU4-P251T. Relative to TrCoV 162275, the identity of HKU4r-BGI-2020 diverges markedly in the S gene and ORF4b and ORF5 genes. Other HKU4-related CoVs exhibit a 75% to 95% identity to HKU4r-BGI-2020, except in the NTD region of the S gene and ORF 4a, 4b and 5 genes where HKU4r-BGI-2020 differs more markedly.

**Fig. 3.**
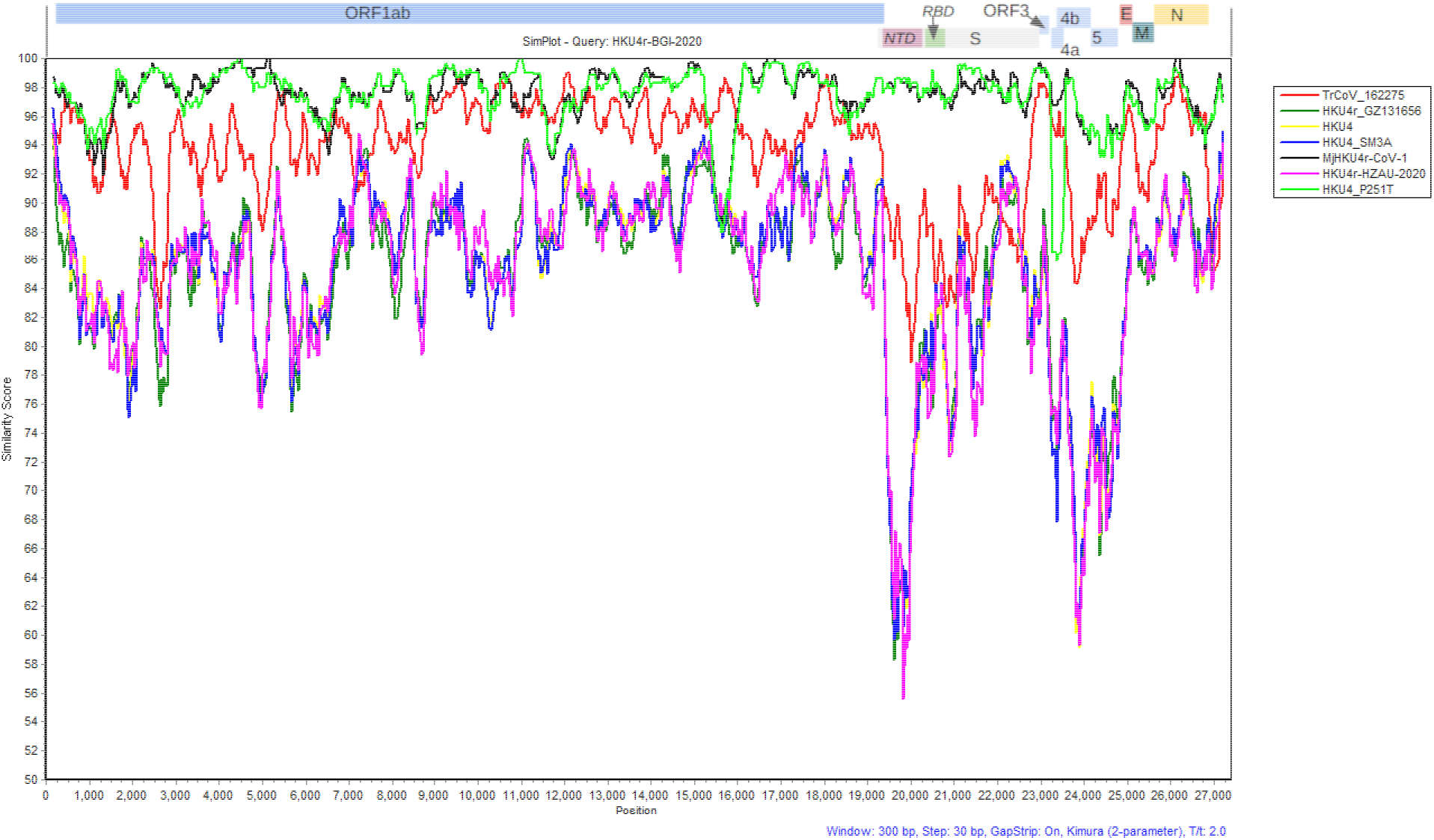
HKU4r-BGI-2020 similarity to seven HKU4-related CoVs. Genomes were aligned using MAFFT. 29 gaps in coverage of HKU4r-BGI-2020 relative to MjHKU4r-CoV-1 and PCoV HKU4-P251T of 50 nt or greater were removed from all sequences after alignment and prior to analysis. Plotted in SimPlot v3.5.1 window size 300 and Step size 30 with otherwise default settings (Kimura 2 parameter model, T/t 2.0, Strip Gaps = True).

### Phylogenetic analysis

Phylogenetic analysis of full merbecovirus genomes using phyML with SMS model selection and aLRT SH-like fast likelihood method shows that HKU4r-BGI-2020 groups with PCoV HKU4-P251T, MjHKU4r-CoV-1 and *Tylonycteris robustula* coronavirus 162275 to form a clade (“clade b” in Fig. 4) distinct from other HKU4-related CoVs which were identified using blast as having highest identity to HKU4r-BGI-2020 (“clade a” in Fig. 4). The HKU4-related CoVs in “clade a”, where known, were sampled from south-western China.

**Fig. 4.**
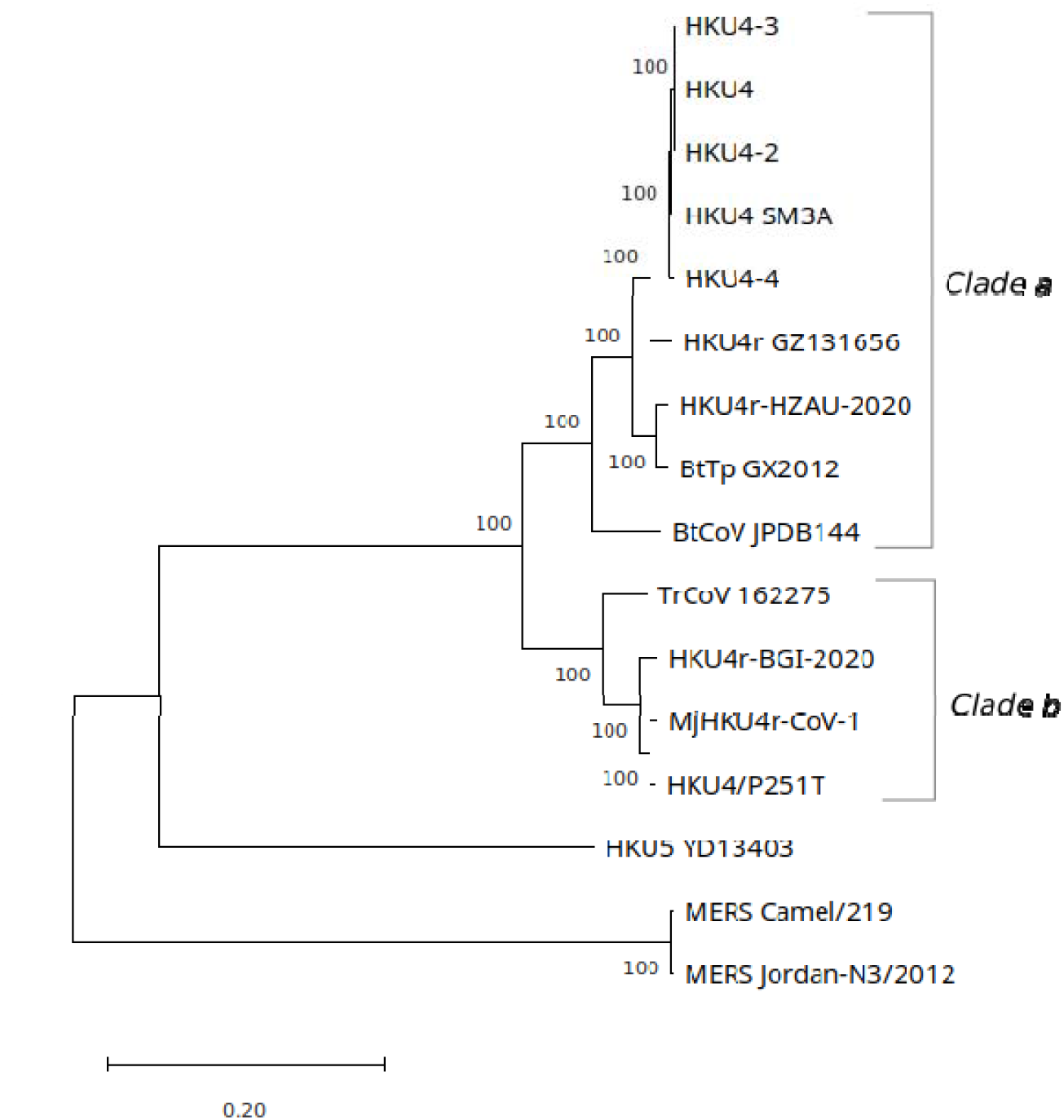
HKU4-related CoV full genome maximum likelihood tree for closest genomes matching HKU4r-BGI-2020, representative HKU4-related CoV genomes from “clade a”, and selected HKU5-related and MERS-CoV genomes. Tree drawn to scale, with branch lengths measured in the number of substitutions per site. Tree rooted on midpoint and displayed using MEGA11.

Phylogenetic analysis of the RdRp gene of Merbecovirus sequences shows a similar relationship as exhibited by the full genome phylogenetic tree (Supp. Fig. 1). HKU4r-BGI-2020 sits in a basal sister relationship with PCoV HKU4-P251T and MjHKU4r-CoV-1. Also, together with *Tylonycteris robustula* coronavirus 162275, it forms a clade separate from other HKU4-related CoVs.

However, more HKU4-related CoV partial RdRp sequences are available compared with full RdRp sequences (Supp. Text). A phylogenetic tree was generated based on a 395-nt fragment of the RdRp gene. Multiple *Tylonycteris robustula* (Greater bamboo bat) CoVs, along with *Pipistrellus coromandra*-hosted PREDICT_CoV-34/KHP13-GT1-0003, group with HKU4r-BGI-2020, PCoV HKU4-P251T, and MjHKU4r-CoV-1 as a distinct clade separate from other HKU4-related CoVs (Fig. 5).

**Fig. 5.**
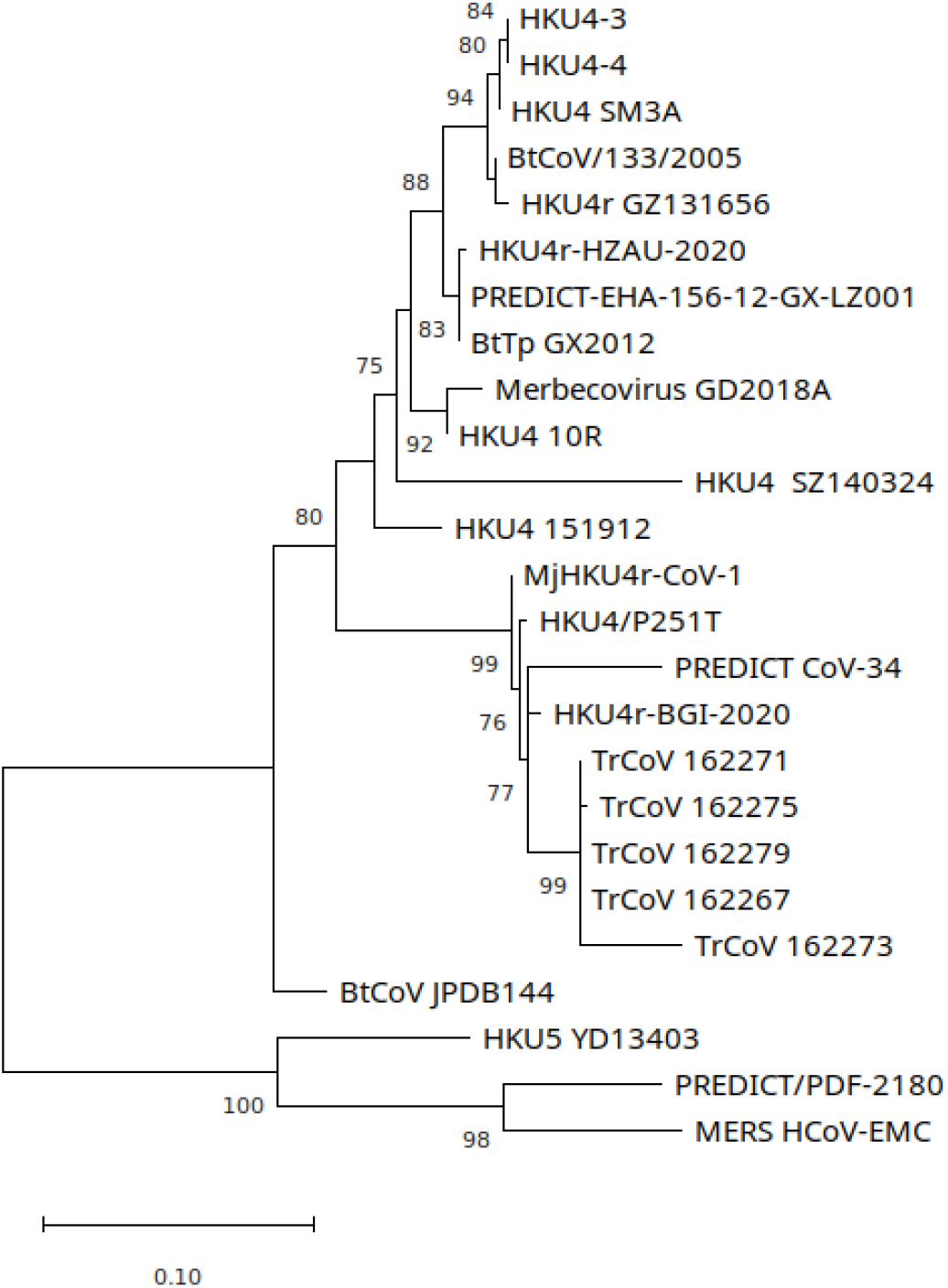
Maximum likelihood tree of a 395-nt partial RdRp coding region. Tree drawn to scale, with branch lengths measured in the number of substitutions per site. Tree rooted on midpoint and displayed using MEGA11.

Both the S gene, S protein and N gene phylogenetic trees also place TrCoV 162275, HKU4r-BGI-2020, MjHKU4r-CoV-1, and PCoV HKU4-P251T in a clade separate from other HKU4-related CoVs (Supp. Figs. 2 to 4). In the ORF3-5 gene region, HKU4r-BGI-2020 is the most evolved of the HKU4-related CoVs in its clade (Supp. Fig. 5).

### Spike Protein

Although the recovery of the receptor binding motif (RBM) sequence of the spike (S) protein is incomplete, we find three amino acid variations are unique to HKU4r-BGI-2020 (Supp. Fig. 6). While 10 amino acids in the RBM of TrCoV 162275 and 17 amino acids in the RBM of HKU4 differ when compared with HKU4r-BGI-2020 (Supp. Fig. 7).

Overall, when missing sections in HKU4r-BGI-2020 are excluded and the gaps ignored, the RBM of HKU4r-BGI-2020 has a 94% amino acid similarity to both MjHKU4r-CoV-1 and PCoV HKU4-P251T, and an 80% similarity to TrCoV 162275. However, the receptor binding domain of HKU4r-BGI-2020 has the highest amino acid similarity to MjHKU4r-CoV-1, at 97%, and an 88% similarity to TrCoV 162275 (Supp. Info. 2.4).

Key sites at the S1/S2 junction and S2’ cleavage sites in HKU4r-BGI-2020 are consistent with related HKU4-related CoVs of the same novel clade. However a region covering a putative furin cleavage motif identified by Chen et al. (2023) in Mj-HKU-CoV-1 (and variants) was not recovered from NGS data (Supp. Text).

A review of evolutionary selection pressure on the spike protein, shows that interestingly, when HKU4r-BGI-2020 is compared against PCoV HKU4-P251T and MjHKU4r-CoV-1, strong purifying selection pressure is evident over the 5’ end of the NTD. However at the 3’ end of the N-terminal domain of the spike protein, while several non-synonymous mutations occur, no synonymous mutations occur, which appears to be unusual (Supp. Text).

### Host analysis

Each SRA dataset in BioProject PRJNA747757 was aligned using minimap2 to the full set of mitochondrial genomes on NCBI downloaded on the 2022-05-22 (Supp. Figs. 15, 16). The most striking finding is the cross-contamination of *Felis catus* and *Manis javanica* genomic material. All *F. catus* datasets contain high coverage of the *M. javanica* mitochondrial genome (at low abundance). Similarly, all *M. javanica* datasets contain high coverage of the domestic cat mitochondrial genome, but with contamination at a significantly higher level. One sample from each of these species further stands out, where lung samples CT_LG (cat lung) and PGN_LG (pangolin lung) have *Capra aegagrus* (wild goat) and *Anser cygnoides* (swan goose) contamination in common.

To further explore the extent of cross-contamination, each SRA data was aligned using 100% identity to a set of all reference mitochondrial genomes from species used in the BioProject and selected species listed in methods. A 60-nt minimum read length filter was then applied to the results. This had the effect of removing the 28-nt F1 direction reads, leaving only R2 100-nt length BGISEQ-500 reads and filtering out short single-end dataset reads. Mitochondrial reads from *C. aegagrus*, *F. catus*, *M. javanica*, and *Sus scrofa* were the most common contaminating species. The *M. javanica* cell datasets were most heavily contaminated with *F. catus* mitochondria, which comprised 42.64%, 6.65%, 14.51%, 15.34%, and 16.32% of all mitochondrial alignments for samples PGN_DM, PGN_LI, PGN_SN, PGN_SH, and PGN_LR, respectively (Supp. Table 1, Supp. Info. 2.5).

Ribosomal RNA (rRNA) from the pangolin cell samples with the highest HKU4-related CoV read counts, PGN_LI, PGN_LG, PGN_ES, along with the *F. catus* liver cell sample CT_LR, *Anolis carolinensis* (green anole) lung cell sample LD_LG, and *Mesocricetus auratus* (golden hamster) lung sample HR_LG were analyzed using a metaxa2 and *de novo* assembly workflow. Local blast against the NCBI nt database used to identify closest homology of the assembled contigs. For sample PGN_LI, only four of the 15 assembled contigs matched *M. javanica* rRNA/DNA or undifferentiated mammalian rRNA/DNA, while 60% of the contigs were bacterial rRNA, while one contig had highest identity to *F. catus* mitochondria. Similarly, for sample PGN_LG, only three of 23 contigs matched *M. javanica* rRNA with abundant bacterial rRNA matches, as well as *A. carolinensis*, *F. catus*, *A. cygnoide*s, *Capra hircus* (domestic goat), *Ambystoma mexicanum* (Salamandra ajolote) mitochondrial matches, and one large contig matching *Mycoplasma fastidiosum* 16S rRNA. Sample PGN_ES rRNA content was again similar, with five of 17 contigs matching *M. javanica* mitochondria of undifferentiated mammalian RNA/DNA, one *M. fastidiosum* 16S rRNA match, and the remaining 65% of contigs comprised of bacterial rRNA (Supp. Data).

Reads from nine *M. javanica* and six *F. catus* single cell SRA datasets in BioProject PRJNA747757 were aligned to full *M. javanica*, *F. catus,* and *C. aegagrus* nuclear genomes using Seal (Bushnell, 2022), with genomic content percentages on the order of mitochondrial genomic content discussed above (Fig. 6).

**Fig. 6.**
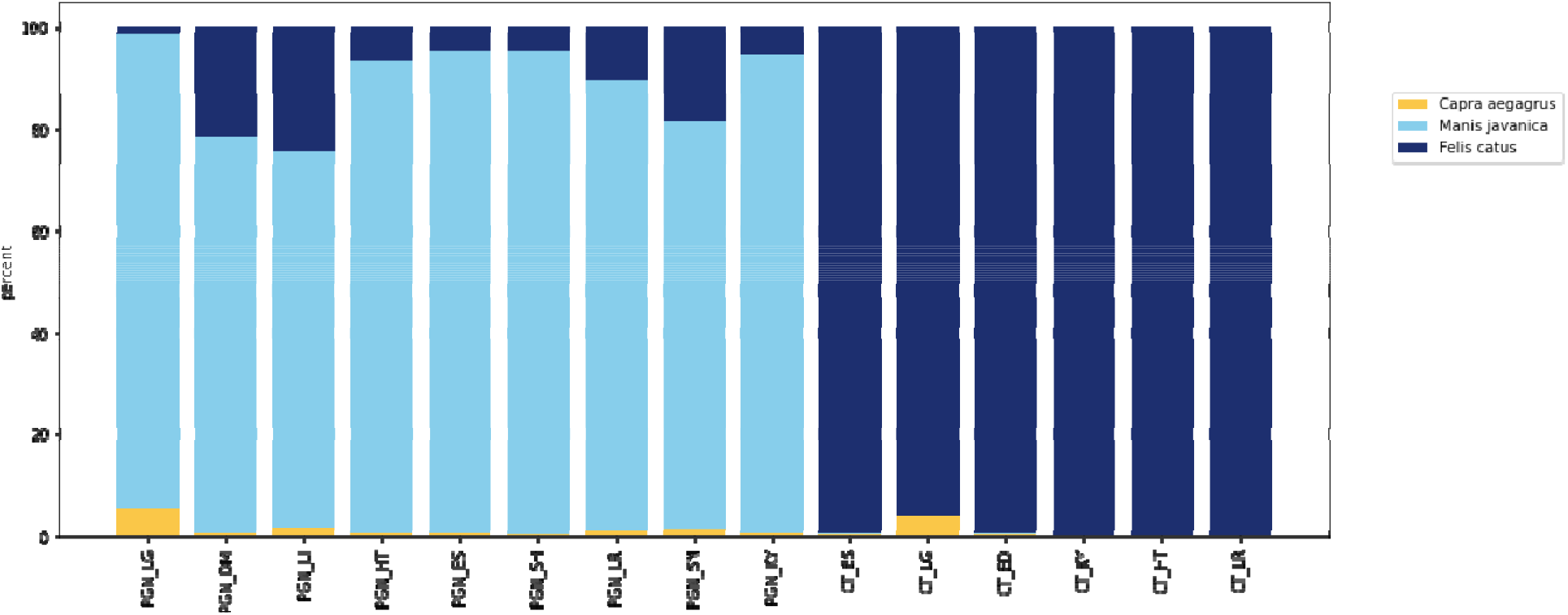
Reads in nine pangolin and six cat SRA datasets in BioProject PRJNA747757 were aligned to full *M. javanica*, *F. catus,* and *C. aegagrus* genomes using seal (Bushnell, 2022) with the “ambiguous = toss” parameter setting.

An analysis of viruses contaminating the NGS datasets is consistent with the mitochondrial and nuclear genome alignment results. In particular Endogenous feline type C virus RD144 matching reads were identified in all pangolin single cell datasets (Supp. Text).

### Host analysis of “clade b” HKU4-related CoVs

While HKU4-related CoVs were first identified in bats in 2006 (Woo et al., 2006), HKU4-related CoVs in pangolin samples were only first documented in August 2022 by Shi et al. (2022). This is over two years after SARS-CoV-2 related CoVs were identified in pangolin RNA-Seq datasets (Liu et al., 2019; Wong et al., 2020; Lam et al., 2020), despite extensive pangolin sampling and sequencing (Lee et al., 2020; Hu et al., 2020; Li et al., 2020; He et al., 2022). Given the cross-contamination of the single cell RNA-Seq datasets in BioProject PRJNA747757 identified here, along with concerns for previous pangolin coronavirus sequencing datasets raised by multiple authors (Hassanin, 2020; Chan and Zhan, 2020; Zhang, 2020; Jones et al., 2021, 2022a, 2022b) we reviewed the RNA-Seq datasets containing these three recently published pangolin-hosted HKU4-related CoVs.

A HKU4-related CoV, HKU4 isolate GX/HKU4-GX/2020 was identified by Cui et al. (2023), in dataset HKU4-GX in BioProject PRJNA901878. This HKU4-related CoV is only two nucleotides different when compared with MjHKU4r-CoV-1 identified by Chen et al. (2023) as discussed previously. We confirmed the presence of CoV HKU4 isolate GX/HKU4-GX/2020 in the dataset, with 14404 mapped reads for 100% coverage and an average read depth of 71 reads (Supp. Fig. 19, Supp. Table 2).

We aligned 86 of the 466 RNA-Seq sequencing datasets in BioProject PRJNA901878 to all mitochondrial genomes on NCBI (Supp. Fig. 20). Datasets for samples HKU4-GX and PPeV-GX were found to be anomalous, in that while they were documented by Cui et al. (2023) as *Manis javanica* anal swab and oropharyngeal swab samples respectively, 99.94% and 99.28% respectively of mitochondrial genome read matching counts where coverage was 10% or greater, were to *Sus* genera (Supp. Info. 4.3, Supp. Figs. 21, 22). Only 338 of the 1,191,655 total mitochondrial genome aligned reads in the dataset for sample HKU4-GX and 993 of the 286,393 total mitochondrial genome aligned reads in the dataset for sample PPeV-GX matched *Manis* genera. On review of the read alignments in HKU4-GX NGS data, after alignment of *de novo* assembled contigs in this dataset against all mitochondrial genomes on NCBI we identified that only 35 of the 338 *Manis* genera mitochondrial aligned reads are potentially valid (Supp. Text).

To confirm taxonomy as indicated by mitochondrial alignment, we undertook full genome reference analysis using three methods: Seal (Bushnell, 2022), XenofilteR (Kluin et al., 2018), and ConcatRef (Jo et al., 2019), and compared results with NCBI STAT analysis. Seal uses alignment free kmer-filtering, and kmer-binning, to split reads into files based on best reference match. In XenofilteR analyses, read mapping statistics were calculated after reads had been separately mapped to two genomes. The ConcatRef workflow analyzed mapping statistics based on best alignment to concatenated genomes. *M. javanica* and *Sus scrofa* genomic content as a percentage of total combined *S. scrofa* and *M. javanica* matching reads were plotted for the three methods (Fig. 7). The *M. javanica* content of datasets PPeV-GX and HKU4-GX calculated using the three methods ranged from 0.034% to 0.31% and was similar to that calculated in *S. scrofa* datasets FL13 and FL16 (0.041% to 0.20%).

**Fig. 7.**
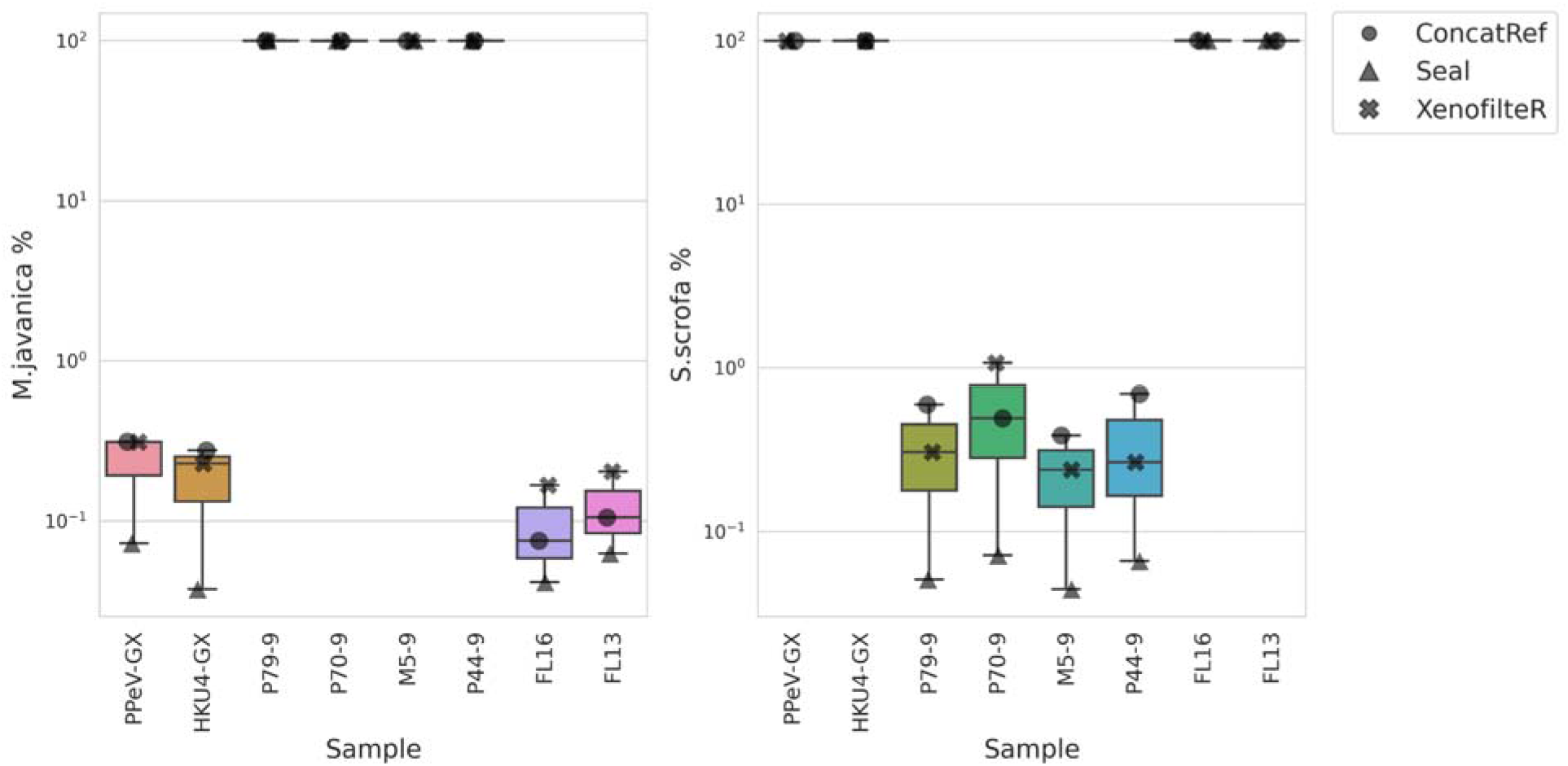
PRJNA901878 selected dataset full genome alignments. Reads in six datasets described as generated from *Manis javanica* samples and four *Sus scrofa* SRA datasets in BioProject PRJNA901878 were aligned to full *S. scrofa*, and *M. javanica* genomes using seal (Bushnell, 2022) with the “ambiguous = toss” parameter setting, ConcatRef and XenofilteR workflows. Percentages shown in log scale.

Analysis of *de novo* assembled contigs using the same three methods gave very similar results (Supp. Fig. 26). Additionally, NCBI STAT taxonomic analysis was consistent with read and contig analysis (Supp. Info. 4.2). Specifically, dataset HKU4-GX was estimated to comprise 60.67% *Sus* genera, 14.18% bacteria, and 4.02% rotavirus. Only 0.02% of the NGS dataset was classified as *Manis* genera.

Given that eight analysis methods, five analyzing reads, three analyzing *de novo* assembled contigs show datasets for samples PPeV-GX and HKU4-GX to contain only trace levels of *Manis* genomic content, we infer with high confidence these two datasets were sequenced from *S. scrofa* origin samples. Sample HKU4-GX with only 35 *Manis* genus (as compared to 1190173 *Sus* genus) mitochondrial genome matching reads, contained 14,404 reads mapping to HKU4 isolate GX/HKU4-GX/2020 for complete coverage (Supp. Fig. 19; Supp. Table 2). We infer that the HKU4 isolate GX/HKU4-GX/2020 viral reads almost certainly could not have been sourced from the low level of *M. javanica* origin content in the sample.

As HKU4 isolate GX/HKU4-GX/2020 was found in a *S. scrofa* sample dataset and not a pangolin sample, we reviewed the NGS datasets of samples containing four close variants of HKU4 isolate GX/HKU4-GX/2020: A96 (MjHKU4r-CoV-1), A97 (MjHKU4r-CoV-2), A98 (MjHKU4r-CoV-3), and A100 (MjHKU4r-CoV-4) sequenced by Chen et al. (2023). We identified that three datasets were sequenced from pangolin samples, and one (A100) from cell culture. Although Chen et al. (2023) only specify the use of Caco-2 cell inoculation for virus isolation, the NGS dataset for sample A100 contains high *Chlorocebus sabaeus* (green monkey) genomic origin content, indicative of Vero cell culture, while the NGS dataset for sample A97 also contains notable *Chlorocebus sabaeus* genomic content (Supp. Text).

Notably, limited evolution between MjHKU4r-CoV-1 and its four variants is indicated by the two or four nucleotide single nucleotide variations (SNVs) between the variants (Supp. Text).

In BioProject PRJNA845961 containing the pangolin CoV HKU4-P251T in the RNA-Seq dataset for sample P251T, we identified a second HKU4-related CoV (“HKU4-P19T-2022”) in the SRA dataset for sample P219T (Supp. Text). The novel CoV is 97% similar to PCoV HKU4-P251T but only 25 reads were recovered for 7% genome coverage. With such a low number of reads the presence of this CoV may be due to contamination from another library or contamination due to index hopping during sequencing.

### Contaminating bacteria

As the RNA-Seq datasets in BioProject PRJNA747757 by Chen et al. (2020, 2021) were sequenced from single cell nuclei libraries bacterial content in the datasets is expected to be only at trace levels and sourced during preparation and sequencing steps. As only 15 of the 29 SRA datasets in BioProject PRJNA747757 were taxonomically classified by NCBI with STAT analysis, we undertook bacterial content analysis using Kraken2 on all samples. We found a Pearson correlation coefficient of 0.86 between NCBI STAT and Kraken2 bacterial percentage contents. One outlier was identified, PGN_KY with a 5.74% bacterial content estimated using NCBI STAT and a 0.1% content using Kraken2 (Supp. Info. 2.2). Samples from only two species exhibit notable variance in bacterial content, *Felis Catus* at 0.2% to 3% and a larger range for *Manis javanica* at between 0.1% and 8% (Fig. 8).

**Fig. 8.**
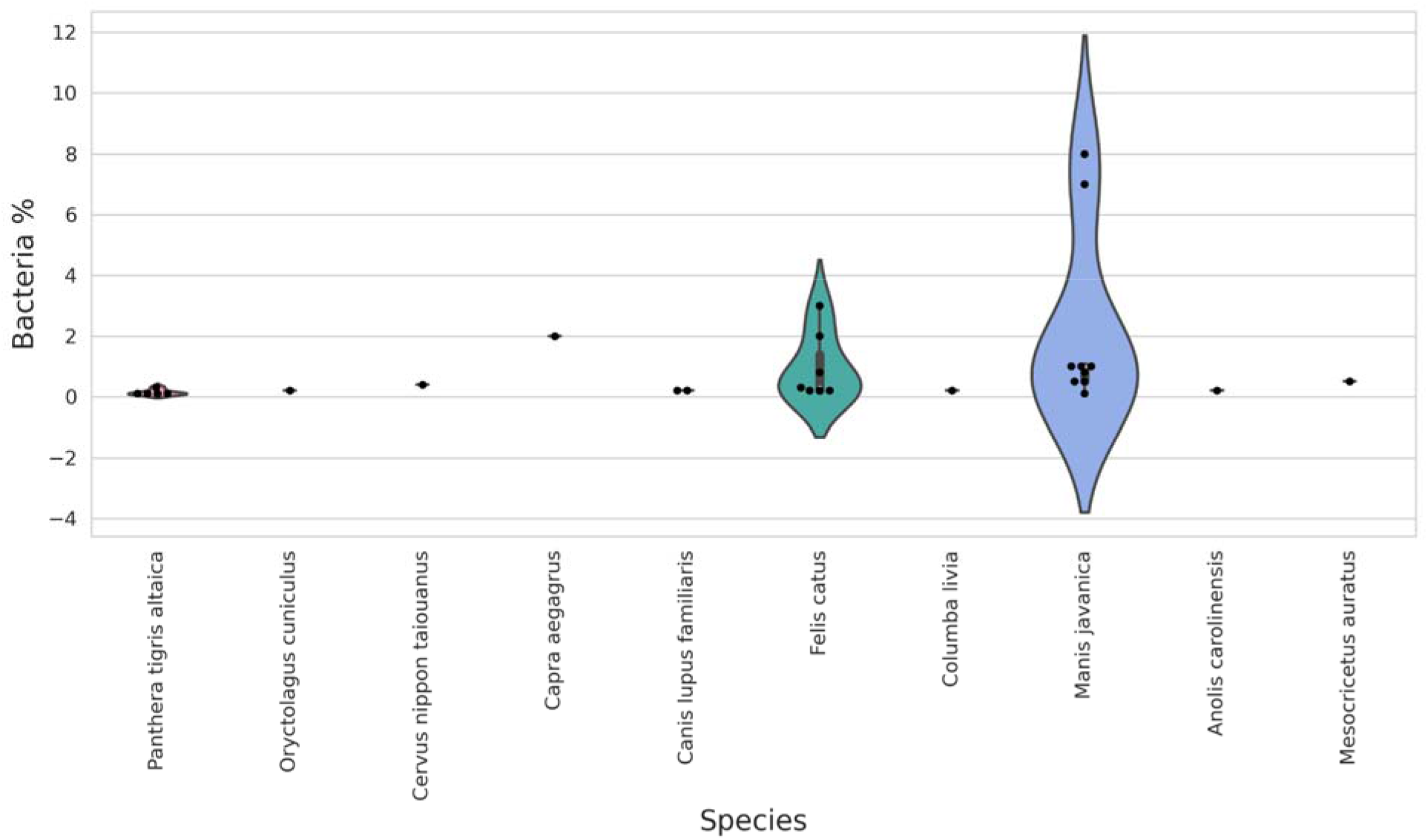
Violin plot of kernel density estimation of bacterial content calculated using Kraken2 for selected SRA datasets in BioProject PRJNA747757, grouped by species. Plotted using seaborn using the “scale=count” parameter. Measured values overlain as black dots.

We note that pangolin large intestine sample PGN_LI, which has the highest bacteria content also has the highest HKU4-related CoV matching read content, with 98% of matching reads in the BioProject. Spearman and Pearson correlation coefficients of 0.5578 and 0.6979 respectively were found between bacteria percentage and HKU4-BGI-2020 read counts in the BioProject, a moderate correlation. An ordinary least squares regression R-squared value between the two series was found to be 0.474, with an F-statistic probability of 2.59e-05 and p-value of 0.000 allowing the rejection of the null hypothesis (Supp. Code).

## Discussion

The novel HKU4r-BGI-2020 coronavirus (CoV) identified here represents the fourth HKU4-related CoV belonging to the newly identified “clade b”, phylogenetically distinct from all previously known HKU4-related CoVs (Fig. 4). HKU4r-BGI-2020 has a 97.34% nucleotide identity to MjHKU4r-CoV-1, a recently discovered pangolin hosted CoV (Chen et al., 2023), 97.23% identity to HKU4-P251T and has a 92.86% identity to the bat *Tylonycteris robustula* CoV isolate 162275.

MjHKU4r-CoV-1 and nearly identical variants MjHKU4r-CoV-2, MjHKU4r-CoV-3, MjHKU4r CoV-4 were identified in four next generation sequencing (NGS) datasets from anal swab samples collected on 2019-11-08 from *Manis javanica* pangolins seized by Guangxi customs, and smuggled from “Southeast Asian countries” (Chen et al., 2023). A fourth variant differing by two nucleotides from MjHKU4r-CoV-1, HKU4 isolate GX/HKU4-GX/2020 (Supp. Fig. 6), was documented as identified in a *M. javanica* pangolin anal swab sample collected in Guangxi province in 2020 (Cui et al., 2023). PCoV HKU4-P251T was detected in a pangolin smuggled into China from inland Asia in 2018–2019, with its mitochondrial genome phylogenetically grouping with Myanmar and Yunnan province pangolins (Shi et al., 2022). It is interesting to note the discovery of a partial HKU4-related CoV “HKU4r_P219T_2022” with a 97.48% identity to PCoV HKU4-P251T had only 25 matching reads in the NGS dataset for sample P219T. With such a low read count it is uncertain if the CoV stems from accidental contamination. The final member of the clade, *Tylonycteris robustula* CoV 162275 was sampled by researchers from the University of Hong Kong in Yunnan province on 2016-08-18.

Surprisingly, we identified that the NGS datasets generated from samples HKU4-GX, PPeV-GX and GX19-89 in BioProject PRJNA901878, are almost entirely comprised of *Sus scrofa* genomic content and not of *M. javanica* (HKU4-GX, PPeV-GX) or *Rhizomys pruinosus* (Bamboo rat) (GX19-89) origin as described by Cui et al. (2023). In sample HKU4-GX, based on the full coverage of HKU4 isolate GX/HKU4-GX/2020 at good read depth, and a very low level of *M. javanica* genomic content in the sample (only 35 reads of the 79.9 million reads in the dataset match the *M. javanica* mitochondrial genome), we infer it implausible that the CoV was related to the presence of *M. javanica* material in the sample. This leads to an important forensic genomic principle that the ratio of virus to animal reads can be used to distinguish whether a particular animal is a likely host. More examples would be necessary to propose a quantitative cut-off for the determination of this general principle.

Pangolin hunnivirus isolate GX/HKU4-GX/2020 was also found to be present in this dataset. This almost certainly indicates that the presence of CoV HKU4 isolate GX/HKU4-GX/2020 in dataset HKU4-GX is not related to infection of the pig from which the sample was taken as this hunnivirus phylogenetically clusters with pangolin hunninviruses GD/P47-7/2019 and pangolin PicoV GD/P70-2/2019. Furthermore, HKU4 isolate GX/HKU4-GX/2020 is nearly identical to pangolin CoV MjHKU4r-CoV-1. The only viable explanation for the presence of HKU4 isolate GX/HKU4-GX/2020 and pangolin hunnivirus isolate GX/HKU4-GX/2020 in the *S. scrofa* NGS dataset is that they are laboratory research related. We do not speculate on how this may have occurred.

The distribution of the MERS-CoV and HKU4-related CoV receptor, dipeptidyl peptidase-4 (DPP4) in tissues has been shown to affect tissue tropism for MERS-CoV and HKU4-related CoVs. In insectivorous common pipistrelle bats, DPP4 expression occurs in intestine and kidney cells, with negligible expression in respiratory tract lining cells, indicating a fecal-oral transmission route (Widagdo et al., 2017). In contrast, in camels, DPP4 is expressed in the upper respiratory tract epithelium, while in humans, DPP4 is expressed in the lower respiratory tract lining and kidneys (Widago et al., 2016). The lack of DPP4 expression in the upper respiratory tract may be the main cause of limited MERS-CoV human-to-human transmission (Meyerholz et al., 2016; Widago et al., 2016). The distribution of DPP4 in pangolin organs was characterized by Chen et al. (2021), who show that DPP4 is expressed in the largest percentage of cells in the kidney, spleen, and lung. DPP4 expression was found in smaller percentages in the heart, liver, and large intestine organs. The duodenum on pangolins was not found to be susceptible to MERS-related CoVs.

The skewed distribution of reads matching the HKU4r-BGI-2020 CoV genome across the pangolin tissue single cell datasets sequenced by Chen et al. (2021) appears inconsistent with this DPP4 expression profile. 98% of reads matching HKU4r-BGI-2020 were found in the pangolin large intestine dataset PGN_LI. When compared on a per-million read basis (322 reads per million reads), this level is 36× of that found in the pangolin esophagus PGN_EU dataset (8.88 reads per million reads). Stomach, liver, lung, heart and duodenum sample datasets had only trace levels of the virus (0.01 to 0.62 reads per million reads). No HKU4-related CoV reads were found in the kidney or spleen sample datasets — two of the organs where DPP4 expression is highest. The lack of presence in organs to which this virus has a natural tropism increases the likelihood these animals were not infected during life but rather the sequences were acquired in the laboratory during processing of organ samples and their subsequent analysis.

Phylogenetically, HKU4-BGI-2020 forms a clade (“clade b” in Fig. 4) with pangolin coronaviruses MjHKU4r-CoV-1 (and near identical HKU4 isolate GX/HKU4-GX/2020), PCoV HKU4-P251T and *Tylonycteris robustula* bat CoV 162275 at the full-genome level, as well as for RdRp, and the S and N genes. HKU4-BGI-2020 is found consistently in a basal sister relation to MjHKU4r-CoV-1, and PCoV HKU4-P251T for the full genome, RdRp, S and N genes — but for the ORF3-5 gene region — is found to be the most evolved of the clade. TrCoV 162275 is found consistently in a basal sister relationship to HKU4-BGI-2020, MjHKU4r-CoV-1 and PCoV HKU4-P251T. It is interesting to note two other recent betacoronavirus discoveries made through forensic analysis of contamination of NGS datasets also exhibit basal sister relationships to closely related CoVs. GX_ZC45r-CoV exhibits a basal sister phylogenetic relationship to SARS-2-related Guangxi pangolin CoVs in the NSP4, NSP10 and partial RdRp regions (Jones et al., 2022b). While HKU4-HZAU-2020, a HKU4-related cDNA clone, again sits in a basal sister relationship to its three most closely related HKU4-related CoVs (Jones et al., 2023).

Pangolins are solitary creatures, except for three-to five-day periods within an approximately three-month mating season (Hua et al., 2015). As such, the potential for coronavirus transmission between pangolins is limited. This contrasts with insectivorous bats where co-roosting between species, and close physical contact is the dominant indicator for inter-species viral spillover (Willoughby et al., 2017; Ruiz-Aravena et al., 2022). For this reason, although pangolins have been proposed as natural coronavirus hosts (Zhang et al., 2020; Peng et al., 2021), this is unlikely. Indeed no CoVs were detected in three hundred and thirty *M. javanica* pangolins in Malaysia sampled between 2009 and 2019 (Lee et al., 2020). Pangolins as intermediate SARS-CoV-2-related coronavirus hosts have been proposed by Lam et al. (2020) and Xiao et al. (2020). Yet both *M. javanica* and *M. pentadactyla* pangolins are ground dwelling, and inhabit grasslands and forests with a diet of ants, termites, and insect larvae (Hua et al., 2015). The potential for the regular close contact with bats that would be required for transmission of coronaviruses over the 2017-2020 period in which pangolins have been identified with beta-coronaviruses is very low. A similar limitation for potential transmission of MERS-CoV from bats to humans was concluded by Meyer et al. (2014). We infer pangolins are also unlikely to be intermediate betacoronavirus hosts. Indeed, the analysis of Chen et al. (2020, 2021) indicated that while some pangolin cell types may be susceptible to MERS-related CoVs, pangolin tissues are poorly susceptible to SARS-related CoVs.

A third hypothesis, that pangolins are incidental hosts, appears to be the only plausible natural infection scenario. Lee et al. (2020) propose the exposure of pangolins to infected humans or animals during wildlife trafficking. Chan and Zhan (2020), noting that Guangdong (GD) pangolin CoV identification and analyses were based on samples from a single batch of pangolins, proposed pangolin infection by other infected animals housed together during smuggling. Choo et al. (2020) and Wenzel (2020) propose the possibility of incidental infection of pangolins by humans via a reverse zoonosis or anthroponosis.

The first submission of a HKU4-related CoV from a pangolin sequencing dataset was PCoV HKU4-P251T on 2022-03-25. Indeed, prior to the Chen et al. (2020), single-cell sequencing study published on 2020-06-14, HKU4-related CoVs had never been identified in any pangolin metagenomic sequencing dataset, or via any testing of pangolins for coronaviruses. This indicates that the infection of pangolins by HKU4-related CoVs is extremely rare. This is consistent with our assessment that pangolins are unlikely to be natural reservoirs or intermediate reservoirs for SARS-related CoVs (Jones et al., 2021; 2022a; 2022b) or HKU4-related CoVs.

Cross-contamination is evident across samples in the BioProject PRJNA747757 by Chen et al. (2020, 2021). Each sample in the BioProject was sequenced on unique flowcell/lane combinations with for example CT_LG and PGN_LG sequenced on the same flowcell but on different lanes. Single cell RNA-Seq is generally conducted on individual cells of a population of the same type. As such multiplexing between samples in this BioProject was unlikely to have been used and we infer that index-hopping associated contamination between samples can be excluded. Contamination upstream of sequencing is thus inferred to be most likely. Low levels of *M. javanica* origin genomic material was found in *F. catus* sequencing datasets, while a higher content of *F. catus* cell contamination of *M. javanica* datasets was found. Pangolin lung (PGN_LG) in particular and cat lung (CT_LG) have similar contamination profiles and we infer pangolin lung tissue origin cells to have contaminated the cat lung dataset.

The origin of the HKU4-related CoV reads in BioProject PRJNA747757 is uncertain and three hypotheses are presented: 1) the sequences are contamination from a source unrelated to this BioProject, 2) the sequences stem from a laboratory infectivity experiment of pangolin cells to a HKU4-related CoV or other laboratory experimentation, or 3) the pangolin was naturally infected with the HKU4-related CoV.

A case for a natural infection can be made given that:

i. of the 11 animal species sequenced by Chen et al. (2020,2021) in 29 RNA-Seq datasets, only pangolin datasets (seven of the nine pangolin cell datasets) were found to contain the novel HKU4-related CoV, but not datasets from any other animal;
ii. the amino acid sequence of the RBD of HKU4r-BGI-2020 is 97% similar to a documented pangolin-hosted CoV, MjHKU4r-CoV-1, and only 88% similar to the closest bat-hosted CoV TrCoV 162275.

However, several observations are inconsistent with a natural infection:

i. the *M. javanica* pangolin sample obtained by Chen et al. (2020, 2021) was collected from a pangolin which died of natural causes at the Guangdong Provincial Wildlife Rescue Center, with presumably no observable signs of a HKU4-related CoV infection, since no evidence of infection was discussed by Chen et al. (2020, 2021);
ii. given the very low prevalence of HKU4-related CoVs in pangolins, which were never documented prior to PCoV HKU4-P251T deposited on GenBank on 2022-03-25, it is exceedingly unlikely that a single pangolin taken at random from the Guangdong Provincial Wildlife Rescue Center would be infected with a novel HKU4-related CoV, and that the infected pangolin cells would be by chance used for MERS-CoV receptor expression analysis;
iii. for bat and human cells, HKU4-CoV uses DPP4 as the receptor (Yang et al., 2014) and if pangolin cells are able to be infected, we infer DPP4 is likely to be the receptor. However, the distribution of reads matching HKU4r-BGI-2020 sequences is not associated with cells from organs expressing the highest amounts of DPP4.
iv. bacterial contamination percentage of pangolin samples shows a moderate correlation to HKU4-related CoV content. Notably, the sample with highest bacterial contamination contains by far the highest HKU4-related CoV read abundance. We note a correlation between bacterial contamination and virus contamination has been previously identified in Wuhan sequenced datasets (Quay et al., 2021a, 2021b).

Two further observations are concerning and further clarification from the dataset authors is required. For the CoV with closest identity to HKU4r-BGI-2020, one of the five near identical variants is found in a *Sus scrofa* metagenomic dataset, with only trace levels of *M. javanica* content. As HKU4 isolate GX/HKU4-GX/2020 is unlikely to be *Sus scrofa*-hosted and is almost certainly not *M. javanica* hosted, its presence in the NGS data is inferred to be laboratory research related. Two other variants contain detectable Vero cell genomic content, a cell line widely used for coronavirus isolation and research (Lam et al., 2020; Kleine-Weber et al., 2018), but the use of Vero cells was not documented by the authors (Chen et al., 2023).

It is unlikely that accidental contamination of the BioProject by unrelated HKU4 CoV research would only impact pangolin cell datasets. It is even less probable that the contamination would be by a coronavirus most closely related to a recently published pangolin-hosted HKU4-related CoVs. As such we infer accidental contamination by unrelated research is an unlikely source for HKU4r-BGI-2020.

We infer that an infectivity experiment using a full-length virus or other laboratory research on the novel HKU4-related CoV is the most likely reason for the presence of HKU4r-BGI-2020 in NGS datasets in the BioProject. We note that an unpublished complete HKU4-related CoV DNA clone was being researched in Wuhan in late 2019 (Jones et al., 2023). Although HKU4r-BGI-2020 is only 85.13% identical to the HKU4r-HZAU-2020 clone and thus is not closely related, HKU4r-HZAU-2020 was only discovered due to accidental contamination. It is plausible that other undocumented HKU4-related CoV research may have been ongoing.

The limitations of this study are that we do not have access to samples, RNA-Seq libraries or detailed laboratory protocols or notebooks to independently verify sequencing results. As a matter of completing the scientific record in light of the importance of these findings, we urge Chen et.al. (2020, 2021) to publicly release their single-cell RNA sequencing dataset of *S. scrofa* cells, as part of their original dataset that was deposited in CNP0001085 (https://db.cngb.org/cnsa/). We also call on Chen et al. (2020, 2021) to review the HKU4-related CoV discovery we report here, clarify the source of the virus in sequenced datasets, and publish the complete genome. We additionally invite Chen et al. (2023) to revise the assertion that HKU4 isolate GX/HKU4-GX/2020 was identified in a pangolin anal swab sample, and clarify the origin of the virus. We further request Shi et al. (2022) to clarify the origin of HKU4-related reads in the RNA-Seq dataset for sample P219T and publish the complete genome if available. Finally we request Cui et al. (2023) to document why the NGS dataset for sample A100 containing MjHKU4r-CoV-4 contains a high proportion of Vero cell genomic content, a cell line not documented in their research, and clarify why Vero cell genomic content was also found in sample MjHKU4r-CoV-2.

## Conclusion

Here we discover a previously undocumented HKU4-related coronavirus (CoV), which phylogenetically groups in a novel clade with three HKU4-related CoVs recently published in 2022 and 2023. The novel CoV was identified in single cell RNA-Seq datasets sourced from a single pangolin which died of natural causes. Given that no HKU4-related CoV had ever been detected in any published pangolin RNA-Seq dataset prior to Chen et al. (2020) and the low rate of detection of HKU4-related CoV in pangolin NGS datasets identified by Shi et al. (2022), Cui et al. (2023) and Chen et al. (2023), it is exceedingly unlikely that a randomly selected wild pangolin, whose organs were used for DPP4 expression analysis, would be naturally infected by a previously undocumented HKU4-related CoV. The bulk of HKU4-related reads, 98%, were found in a large intestine single cell nuclei RNA-Seq dataset. Of all 29 single cell RNA-Seq datasets, this dataset contained the highest bacterial contamination of approximately 8%, while overall a moderate correlation between bacterial content and viral read content was found. Additionally, while large intestine cells were found by the authors to exhibit low expression of pangolin DPP4, kidney and spleen single cell datasets, which exhibit highest DPP4 expression, did not contain any detectable HKU4-related CoV matching reads. For these reasons we conclude that the novel HKU4-related CoV is likely laboratory research-related and not related to a natural infection of pangolin cells in which the CoV was identified. We further identify that the RNA-Seq dataset containing HKU4 isolate GX/HKU4-GX/2020 is a *Sus scrofa* metagenomic dataset and not a *Manis javanica* pangolin dataset as documented. Two other close variants of this isolate, differing by only two nucleotides, MjHKU4r-CoV-2 and MjHKU4r-CoV-4, were found in NGS datasets containing a minimum Vero cell content of 3% and 30% of mammalian genomic content respectively. Our findings raise significant concerns for the veracity of claims of natural pangolin infection by HKU4-related CoVs and we call on the authors of this research to respond to our concerns through addendums or corrections to their published papers.

## Methods

NCBI STAT analysis (Agarwala et al., 2018; Katz et al., 2021) and was conducted on each SRA dataset in BioProject PRJNA747757 and selected SRA datasets in BioProjects PRJNA901878 and PRJNA845961 (Supp. Info. 2, 4, 6).

All SRA datasets analyzed were pre-processed using fastp with the “--detect_adapter_for_pe” parameter setting for paired end datasets and default settings for single end fastq files. Microbial analysis using Fastv (Chen S. et al., 2020) against the Opengene vial genome kmer collection “microbial.kc.fasta.gz” (https://github.com/OpenGene/UniqueKMER) downloaded on 2020-06-11 was conducted for all analyzed SRA datasets.

All read alignments were conducted using Minimap2 version 2.24 (Li et al., 2018) with the following parameters “-MD -c -eqx -x sr --sam-hit-only -- secondary=no -t 32” unless otherwise specified. Nucleotide to protein translation was conducted using the Expasy translate tool (https://web.expasy.org/translate/).

For Bioproject PRJNA747757, reads from sample datasets PGN_LI, PGN_LG and PGN_ES were pooled and *de novo* assembled using MEGAHIT v1.2.9 (Li et al., 2015) with default settings. We then aligned both the pooled reads from PGN_LI, PGN_LG and PGN_ES to PCoV HKU4-P251T using minimap2. A preliminary consensus genome was generated from aligned reads using samtools consensus with default settings. Gaps in the preliminary consensus genome were filled using the PCoV HKU4-P251T genome. All SRA datasets in BioProject PRJNA747757 were then aligned using minimap2 to this infilled genome. Reads aligning to the infilled consensus genome were found in seven pangolin cell datasets, and were concatenated into a single BAM file using samtools merge. A consensus genome for HKU4r-BGI-2020 was then generated using samtools and ivar (Grubaugh et al., 2018) as per the bash script “samtools_ivar_consensus.sh” in supplied code. The final HKU4r-BGI-2020 genome is supplied in Supp. Data.

### Phylogenetic Trees

All multiple sequence alignment was conducted using MAFFT v7.490 (Katoh and Standley, 2013) using the “--auto” setting, and trimmed using UGENE v46.0 (Okonechnikov et al., 2012). Phylogenetic analysis of aligned and trimmed sequences was conducted using the following workflow, unless otherwise indicated: Fasta files were converted to phylip format using seqmagick v0.8.4 (https://github.com/fhcrc/seqmagick). Phylip files were then analyzed using PhyML v3.0 (Guindon et al., 2010) online (http://www.atgc-montpellier.fr/phyml/) with smart model selection (SMS) (Lefort et al., 2017) model selection and aLRT SH-like fast likelihood method. Trees were saved as newick format and imported into MEGA11 (Tamura et al., 2021) or iTOL (Letunic and Bork, 2021) for visualization. Bootstrap support values less than 70% were not displayed. Trees are displayed rooted on midpoint unless otherwise indicated.

To generate a maximum likelihood tree for full genomes for MjHKU4r-CoV-1, its four close variants and *Tylonycteris robustula* CoV 162275, model selection was conducted using MEGA11. The GTR+I model was found to have lowest bayesian information criterion and a maximum likelihood tree was generated using 100 bootstrap replicates.

### Host analysis

All fastp processed SRA datasets in BioProjects PRJNA901878 and PRJCA002517 and selected SRA datasets in BioProjects PRJNA901878 were aligned to all mitochondrial genomes on NCBI downloaded on 2022-05-05, using minimap2 with parameters specified above. Samtools sort and Samtools coverage were used to calculate mapping statistics.

Mitochondrial alignment using 100% identity for each SRA dataset in BioProject PRJNA747757 was generated using bowtie2 v2.4.2 with the following parameter setting: “--score-min L,0,0”. The set of mitochondrial genomes used consisted of the following accession codes: MN816163.1, KY018919.1, MG196309.1, MT335859.1, NC_012920.1, NC_007393.1, KP257597.1, KX355640.1, NC_053269.1, MK089295.1, MK251046.1, NC_001700.1, NC_028161.1, NC_009684.1, HM185182.2, NC_013978.1, EU747728.2, NC_013276.1, KX467593.1, GU684806.1, NC_007936.1.

#### Seal

Seal is a k-mer-matching tool in the BBTools suite (Bushnell, 2022) which allows alignment-free, kmers matching to multiple reference sequences. We used BBMap version 38.87 to run seal with alignments to either two or three nuclear genomes using the “ambiguous = toss” parameter setting. Code is provided in the Seal_kmer_match.ipynb jupyter notebook. Plot generation code is provided in Plots_Seal_Stats.ipynb.

#### ConcatRef

A ConcatRef workflow (Jo et al., 2019) was implemented whereby fasta files for multiple nuclear genomes were concatenated and reads or contigs aligned using minimap2 (Alignment_minimap2_raw_reads.ipynb, Alignment_minimap2_contigs.ipynb). Mapping statistics were than calculated using the jupyter notebook ConcatRef_Stats.ipynb and the bash script concatref_sam_proc.sh

#### Disambiguate

A disambiguate workflow (Jo et al., 2019) was run using cvbio v3.0.0 (Valentine, 2020). Fastp processed SRA datasets in BioProject PRJCA002517 were aligned to three full length genomes in serial, GCF_015252025.1_Vero_WHO_p1.0_genomic.fna, GCF_000001405.39_GRCh38.p13_genomic.fna, and GCF_014570535.1_YNU_ManJav_2.0_genomic.fna using bwa-mem v0.7.17 (Li, 2013) using default parameters (Alignment_bwamem2_raw_reads.ipynb). Cvbio was used to disambiguate read mappings (Cvbio_Disambiguate.ipynb).

#### XenofilteR

XenofilteR v1.6 (Kluin et al., 2018) was used to filter reads and mapped separately to two nuclear genomes as per code in XenofiltR_script.R and XenofiltR_contigs_script.R.

#### Metaxa2

Reads were classified using metaxa2 against a local copy of the nt database using the workflow in metaxa2_paired.sh. De novo assembled contigs were then classified against a local copy of the nt database using the Blast_nt_contigs.ipynb jupyter notebook.

### Selection pressure analysis

MACSE V2.07 (Ranwez et al., 2018) was used to codon align spike gene sequences using the following command “java -jar macse.jar -prog alignSequences -seq spike_sequences.fa”. SNAP v2.1.1 (Korber, 2002) was used to generate dN/dS plots.

### Contaminating virus analysis

All SRA datasets in BioProjects PRJNA747757, PRJNA901878 and PRJNA845961 were aligned using minimap2 to a NCBI viral dataset downloaded on 2021-11-28, concatenated with all coronaviruses on NCBI downloaded on 2022-12-26. Viruses with coverage greater than 10% were then shortlisted and compared with fastv microbial analysis. SRA datasets were then re-aligned to the shortlisted virus set using minimap2. We additionally aligned all SRA datasets to a set of SARS-related and HKU4-related gomes. Notably we detected a problem with the PCoV HKU4-P251T gnome (OM009282.1) whereby the first 75nt of the genome is clearly in error. We replaced the first 75nt with a 73nt sequence at the 5’ end of the genome generated from *de novo* assembly of HKU4r-BGI-2020, which we term OM009282r. Selected results were plotted as per code in Plots_heatmaps_viruses_PRJNA747757.ipynb.

### Contaminating bacterial analysis

Reads were classified using Kraken2 v2.1.1 (Wood et al., 2019) against the NCBI standard database consisting of Refeq archaea, bacteria, viral, plasmid, human1, UniVec_Core compiled on 2023-03-14. Krona v2.0 (Ondov et al., 2011) was used for display with plots generated using the following commands:

kraken2 --db k2_standard --paired --classified-out k2_standard#.fq sra_1_fastp.fq sra_2_fastp.fq --output k2_standard.txt --threads 32

cat k2_standard.txt | cut -f 2,3 > k2_standard.krona

ktImportTaxonomy -tax k2_standard_20210517/taxonomy k2_standard.krona.

Statistical correlation between NCBI STAT and kraken2 bacterial taxonomic classification was generated as per code in Plots_bacteria_PRJNA747757.ipynb.

## Supplementary Figures and Tables

Supplementary figures and tables have been deposited on Zenodo at:

doi:10.5281/zenodo.8051589

https://zenodo.org/record/8051589

## Supplementary Text

Supplementary text has been deposited on Zenodo at:

doi:10.5281/zenodo.8051589

https://zenodo.org/record/8051589

## Supplementary Data

Supplementary data including the HKU4r-BGI-2020 genome and metaxa2 rRNA analysis results has been deposited on Zenodo at:

doi:10.5281/zenodo.8051589

https://zenodo.org/record/8051589

## Code

All code can be found at: https://github.com/bioscienceresearch/Novel_HKU4r-CoV_single_cell

