## Supplementary Text for "Identification of a novel HKU4-related coronavirus in single-cell datasets and clade viral host analysis"

^1^ Independent Bioinformatics Researcher, Melbourne, Australia; ORCID 0000-0001-7355-0052

^3^ Department of Chemistry and Physics, Halmos College of Arts and Sciences, Nova Southeastern University, Ft. Lauderdale, FL, USA, ORCID 0000-0001-8861-0898

^5^ Youthereum Genetics Inc., Toronto, Ontario, Canada; ORCID 0000-0002-3397-5811

^6^ Atossa Therapeutics, Inc., Seattle, WA USA; ORCID 0000-0002-0363-7651

### Supplementary Text

#### HKU4-related coronaviruses with only partial RdRp sequences

For a 395-nt partial RdRp region with more extensive HKU4-related CoV sampling, multiple *Tylonycteris robustula* coronaviruses and *Pipistrellus coromandra*-hosted PREDICT_CoV-34/KHP13-GT1-0003 group in the same clade as MjHKU4r-CoV-1, PCoV PCoV HKU4-P251T and HKU4-BGI-2020 (Fig. 5). Metadata for TrCoVs 162267, 162271, 162275, and 162279 show that these were sampled by Libiao Zhang from the Guangdong Key Laboratory of Animal Conservation and Resource Utilization on 2016-08-18 from a location in Yunnan province, China. *Pipistrellus coromandra*-hosted PREDICT_CoV-34/KHP13-GT1-0003 was sampled by Lacroix et al. (2017) in Cambodia in 2013 and deposited on Genbank on 2019-02-26. *Tylonycteris robustula* (Greater flat‐headed bats), as well as *Tylonycteris pachypus* (Lesser flat‐headed bats) are both found in bamboo forest, where they form small colonies in hollow bamboo internodes (Liang et al., 2019). The *Pipistrellus coromandra* (Indian pipistrelle) bats sampled by Lacroix et al. (2017) were located in deforested and agricultural regions.

*T. robustula* CoV 162275 was submitted to the NCBI by Liang, J. from the Guangdong Key Laboratory of Animal Conservation and Resource Utilization, Institute of Zoology, Guangdong Academy of Sciences, to which has conducted widespread bat betacoronavirus and pangolin betacoronavirus research (Liu et al., 2019; Liu et al., 2020; Li et al., 2021) .

#### Spike protein

A putative furin cleavage motif RQQR was observed by Chen et al. (2023) in pangolin CoV MjHKU4r-CoV-1 in the S1 region of the S protein, downstream of the RBD. The same motif is found in two other CoVs, the nearly identical GX/HKU-GX/2020 and in PCoV HKU4-P251T. The *Tylonycteris robustula* bat CoV 162275 and MERS-CoV also possess this sequence. However, reads covering this part of the genome were not recovered for HKU4r-BGI-2020 (Supp. Fig. 8).

MERS-CoV possesses a furin recognition motif RSVR|S at the S1/S2 boundary (Kleine-Weber et al., 2018), with the role of furin for mediating MERS-CoV cell entry debated (Matsuyama et al., 2018). Most HKU4-related CoVs possess a STFR|S or SLFR|S sequence at the S1/S2 boundary. However, HKU4r-BGI-2020, PCoV HKU4-P251T, PCoV MjHKU4r-CoV-1 (and CoV GX/HKU4-GX/2020) as well as *Tylonycteris robustula* bat CoV 162275 differ with a double threonine TTFR|S sequence (Supp. Fig. 9). Also, near the S1/S2 boundary where MERS has a human endosomal cysteine protease recognition motif AFN|H, the viruses HKU4r-BGI-2020, PCoV MjHKU4r-CoV-1, and PCoV HKU4-P251T possess a NHT sequence in common with HKU4-related CoVs, but differ in possessing a trailing alanine rather than a serine.

In MERS-CoV, the S2’ cleavage site RSAR is located upstream of the putative fusion peptide in the S2 subunit. The S2’ site, but not necessarily the S1/S2 site, was found to be required for human cell entry (Kleine-Weber et al., 2018) (Supp. Fig. 10). In contrast HKU4-related CoVs possess only a single arginine at this site and are unable to efficiently utilize endogenous human proteases for cell entry (Yang et al., 2014; Wang et al., 2014; Yang et al., 2015). However, Ty-BatCoV HKU4 SM3A (Supp. Fig. 10) has been demonstrated to be able to infect human cells and hDPP4-transgenic mice (Lau et al., 2021).

A notable difference of HKU4r-BGI-2020, PCoV HKU4-P251T, PCoV MjHKU4r-CoV-,1 and *Tylonycteris robustula* bat CoV 162275 as opposed to HKU4-related CoVs where the S gene has been sequenced, is that they possess a three-amino-acid insert relative to HKU4-related CoVs in the NTD of the S protein at an external position (Supp. Figs. 11, 12).

#### Selection pressure

To analyze evolutionary selection pressure, the evolutionary rate ratio of numbers of non synonymous mutations to synonymous mutations (dN/dS) across the spike proteins of HKU4r-BGI-2020 and the closest known HKU4-related CoVs MjHKU4r-CoV-1, PCoV HKU4-P251T, and TrCoV 162275. Positive selection occurs where dN/dS>1, while purifying selection occurs where dN/dS<1 (Jeffares et al., 2015). When each of HKU4r-BGI-2020, MjHKU4r-CoV-1, and PCoV HKU4-P251T are compared with TrCoV 162275, an absence of selection (dN/dS ~ 1) in the N-terminal domain (NTD) region is observed (Supp. Fig. 13). Purifying selection pressure is observed in both the RDB and S2 regions indicating more stringent evolutionary control.

When the spike protein of HKU4r-BGI-2020 is compared with the spikes from PCoV HKU4-P251T and MjHKU4r-CoV-1, strong purifying selection pressure is evident in the upstream half of the N-terminal domain (NTD) (Supp. Fig. 14). Curiously, just upstream, over the 3’ end of the NTD, no synonymous mutations occur in a region with several non-synonymous mutations. Similar relationships are observed over the RBD region.

When the spike proteins from PCoV HKU4-P251T and MjHKU4r-CoV-1 are compared, extreme purifying selection pressure (dN/dS = 0) is found over the entire NTD and 5’ end of the RBD. The complete lack of non-synonymous mutations for a 440 amino acid section, and the low number of synonymous mutations for this region observed between spike proteins for PCoV HKU4-P251T and MjHKU4r-CoV-1, could indicate recent recombination. In a natural recombination scenario, this may indicate PCoV HKU4-P251T and MjHKU4r-CoV-1 host population mixing.

#### Contaminating Viruses

Viruses in datasets in BioProject PRJNA747757 were first identified using NCBI STAT krona analysis and fastv, and a reference set of virus genomes was generated containing viruses with significant coverage. Pangolin CoV HKU4-P251T was added to the reference set. All pangolin cell datasets, as well as seven cat (*Felis catus*), one duck (*Anas platyrhynchos*), one golden hamster (*Mesocricetus auratus*), and one rock dove (*Columba livia*) cell datasets were aligned to this reference set of viruses. Read counts and virus coverage of greater than 10% in the datasets were reviewed (Supp. Figs. 17, 18).

Endogenous feline type C virus RD144 was found at >10% coverage in all nine pangolin cell datasets, but no obvious correlation with PCoV HKU4-P251T (a proxy for HKU4r-BGI-2020 content) was evident. The pangolin lung dataset PGN_LG exhibits the highest non pangolin-hosted virus contamination of the pangolin single cell datasets reviewed. At the same time, cat lung sample CT_LG exhibited the most significant contamination from non cat-hosted viruses of the *felis catus* single-cell samples.

#### Host analysis of “clade b” HKU4-related CoVs

##### GX/HKU4-GX/2020 CoV

After *de novo* assembly of reads in the dataset for sample HKU4-GX and alignment of contigs to all mitochondrial genomes on NCBI, the 5’ end of a 6531-nt contig was found to match the *Manis javanica* isolate T298 mitochondrial genome (Supp. Fig. 23). Blastn analysis of the contig exhibits a best match to *Phacochoerus africanus* spindlin 1 (SPIN1), transcript variant X1, mRNA with a 98.06% identity for 4118nt. An anomalously high read depth was found at this location in both HKU4-GX and PPeV-GX datasets (Supp. Fig. 23). We infer that reads aligning to the *M. javanica* mitochondrial genome between positions 1350 and 1950-nt are spurious and that only 35 of the aligned reads are potentially valid *M. javanica* alignments. On removal of likely invalid alignments, mitochondrial genome coverage was 17.47%. In comparison, the coverage of the *Sus scrofa taiwanensis* mitochondrial genome was 100% by 56858 reads at an average read depth of 515.49 (Supp. Fig. 24).

A third anomalous sample in BioProject PRJNA901878 was also identified, *Rhizomys pruinosus* liver sample GX19-89 (SRR22936497). NCBI STAT taxonomic classification of DZ1, the only other *R. pruinosus* sample in the project, indicates the sampled dataset is of the order Rodentia with low levels of Homo sapiens contamination. However sample GX19-89 is classified as belonging to the Suidae family, with no Rodentia genomic matches identified. Mitochondrial genome alignment shows 7.4x the number of reads aligning to Sus sp. Mitochondrial genomes than to *Rhizomys sp.* (Supp. Info 4.3). Mitochondrial genome analysis of the dataset for sample DZ1 did not contain any *Sus* mitochondrial genome coverage of 10% or more, a minimum cutoff applied (Supp. Info 4.3). Full genome alignments of the NGS dataset for sample GX19-89 using Seal, ConcatRef and XenofilteR workflows indicate the dataset is comprised of a mammalian genomic content of 1-3% *R. pruinosus* and 97-99% *S. scrofa* (Supp. Fig. 25). This is concerning as a novel CoV, Bamboo rat CoV GX/GX19-89/2019, found in the dataset, is proposed to have a Bamboo rat host (Cui et al., 2023).

##### MjHKU4r-CoV-1

We aligned reads in the four SRA datasets containing CoVs MjHKU4r-CoV-1-4 in BioProject PRJCA002517 to a complete NCBI mitochondrial genome reference set (Supp. Figs. 27, 28). Sequencing dataset CRR477154 of sample A96 containing MjHKU4r-CoV-1 contains only *Manis* genera matching mitochondrial sequences at greater than ten percent coverage. However datasets CRR477155-7 from samples A97, A98, and A100 containing HKU4-related CoVs MjHKU4r-CoV-2, MjHKU4r-CoV-3, MjHKU4r-CoV-4 all contain a significant amount of *Homo sapiens* and *Chlorocebus sabaeus* (African green monkey) mitochondrial genome matching reads. To ascertain the mapping statistics of HKU4-related CoVs in the NGS datasets, we aligned each dataset to the CoV present (Supp. Table. 3)

Seal (Bushnell, 2022), ConcatRef (Jo et al., 2019), and Disambiguate using cvbio (Valentine, 2020) workflows were run on reads and *de novo* assembled contigs to determine mapping statistics to the *Manis javanica*, *H. sapiens*, and *C. sabaeus* genomes. Cvbio implements a modified version of Disambiguate (Ahdesmäki et al., 2017) to allow disambiguation of alignments to more than two genomes.

The NGS dataset for sample A100 (CRR477157), of reads unambiguously mapping to these genomes, contains dominantly *H. sapiens* and *C. sabaeus* and only 3.5% to 33.9% *M. javanica*, genomic material (Supp. Fig. 29). Aligned contigs show a slightly higher number of contigs matching *C. sabaeus* over the *H. sapiens* genome (Supp. Fig. 30). In the NGS dataset for sample A97, the lowest *C. sabaeus* content of reads mapping to the three mammalian species identified as present in the BioProject was 3%.

Taxonomic classification of the NGS dataset for sample A100 (CRR477157) using Metaxa2 to identify and classify rRNA was undertaken and results were de novo assembled. 55 contigs were generated and classified against a local copy of the nt database using blastn. While matches were dominantly to bacteria, four *H. sapiens*, two Chlorocebus sp., two Pan sp. and two Manis sp. contigs were identified (Supp. Data). While Metaxa2 classification workflow for NGS dataset for sample A100 (CRR477157) resulted in 39 contigs. Of these, one 648-nt contig best matched a *H. sapiens* mitochondrial genome, one 1182-nt length contig best matched the *Cercopithecus aethiops sabaeus* (*Chlorocebus sabaeus*) mitochondrion. This confirms the presence of both *H. sapiens and C. sabaeus* genomic content in both samples.

Two RNA-Seq methodologies were discussed by Chen et al. (2023). 1) Unpurified RNA of anal swab samples sequenced on the MGI DNBSEQ-T7 platform; 2) sequencing of culture supernatants from anal swab sample-inoculated Caco-2 cells, sequenced on the BGI MGISEQ2000 platform. Fastq headers show instrument ids on NGS datasets for samples A96, A97 and A100 to have the same instrument id. While sample samples A95 was sequenced on a different instrument. Library construction details specified on GSA (Genome Sequence Archive) accession number: CRA006906 specify that library construction for samples A95, A96, A97 and A100 involved inoculation on cells and RNA extraction from supernatant, all sequenced on the MGISEQ-2000RS platform. As NGS datasets for samples A95, A96, A97, of mammalian genome content, contain dominantly *M. javanica* genome aligning reads, we infer the library construction for these NGS datasets was using unpurified RNA as per method 1) above. Library construction for sample A100, which contains dominantly *H. sapiens* and *C. sabaeus* we infer was constructed using method 2. Neither method explains why *C. sabaeus* genomic origin content is found in NGS datasets A97 and A100.

##### MjHKU4r-CoV-1 variants

MjHKU4r-CoV-1, MjHKU4r-CoV-2, MjHKU4r-CoV-4 and HKU4 isolate GX/HKU-GX/2020 all differ by two nt to each other, while MjHKU4r-CoV-3 differs by four nt, and all represent variants of a single strain (Supp. Fig. 31; Supp. Info. 5.4). HKU4r-BGI-2020 contains consensus T58, C3181, T10282, A11905 and T21773 codings, but contains the SNV T11030C relative to MjHKU4r-CoV-1 (Supp. Info. 5.4). Position 20558 in the HKU4r-BGI-2020 genome was not recovered. Interestingly, position 20545 in HKU4 isolate GX/HKU-GX/2020 (position 20558 relative to MjHKU4r-CoV-1), is transitional between cytosine, with a read depth of 23 in supporting NGS data, and thymine with a read depth of 31 reads.

##### PCoV HKU4-P251T

We aligned all SRA datasets in BioProject PRJNA845961 to a NCBI viral dataset, concatenated with all coronaviruses on NCBI. We additionally aligned all SRA datasets to a set of SARS-related and HKU4-related CoVs. We identified 412362 reads in the NGS dataset for sample P251T mapping to a slightly modified version of the PCoV HKU4-P251T with a fix to a problematic 5’ end of the genome as per methods (Supp. Info 6.2). We also detected 25 reads for a 7% coverage of PCoV HKU4-P251T in the NGS dataset for sample P219T (Supp. Info 6.2; Supp. Data). We then aligned both the NGS datasets for samples P251T and P219T to a set of all mitochondrial genomes on NCBI. The dataset for sample P251T was found to contain Homo sapiens mitochondrial contamination, while only Manis genera mitochondrial genome alignments with a greater than 10% coverage were found in the NGS dataset for sample P219T (Supp. Info 6.2).

Interestingly six other NGS datasets in this BioProject had reads matching PCoV HKU4-P251T, but at an even lower level than the dataset for P219T, at between two to six reads. Five datasets had zero single nucleotide variations (SNVs) relative to PCoV HKU4-P251T, one dataset had four SNVs (Supp. Info. 6.2). We infer index hopping could be the cause of the contamination.
