## Supplementary Figures for "Identification of a novel HKU4-related coronavirus in single-cell datasets and clade viral host analysis"

^1^ Independent Bioinformatics Researcher, Melbourne, Australia; ORCID 0000-0001-7355-0052

^3^ Department of Chemistry and Physics, Halmos College of Arts and Sciences, Nova Southeastern University, Ft. Lauderdale, FL, USA; ORCID 0000-0001-8861-0898

^5^ Youthereum Genetics Inc., Toronto, Ontario, Canada; ORCID 0000-0002-3397-5811

^6^ Atossa Therapeutics, Inc., Seattle, WA USA; ORCID 0000-0002-0363-7651

### Supplementary Figures and Tables

#### Phylogenetic Analysis

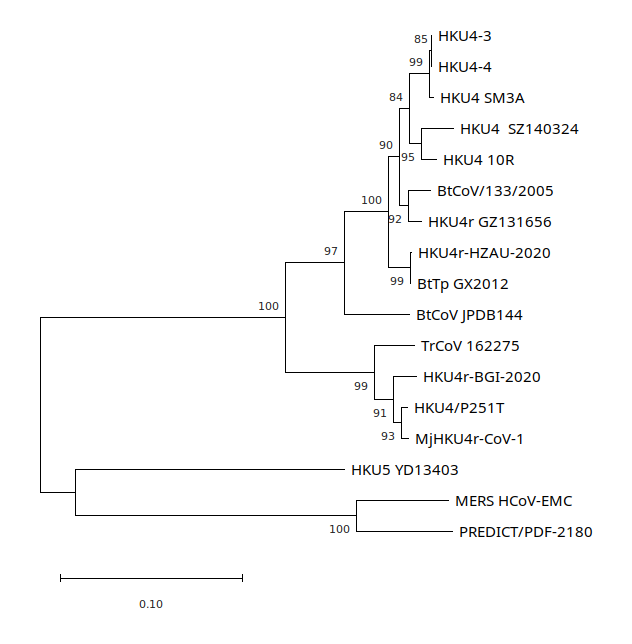

Supp. Fig. 1. HKU4-related CoV RdRp maximum likelihood tree. Tree drawn to scale, with branch lengths measured in the number of substitutions per site, bootstrap values >70% are shown. Tree rooted on midpoint and displayed using MEGA11. Accession numbers and names in Supp. Info. 3.

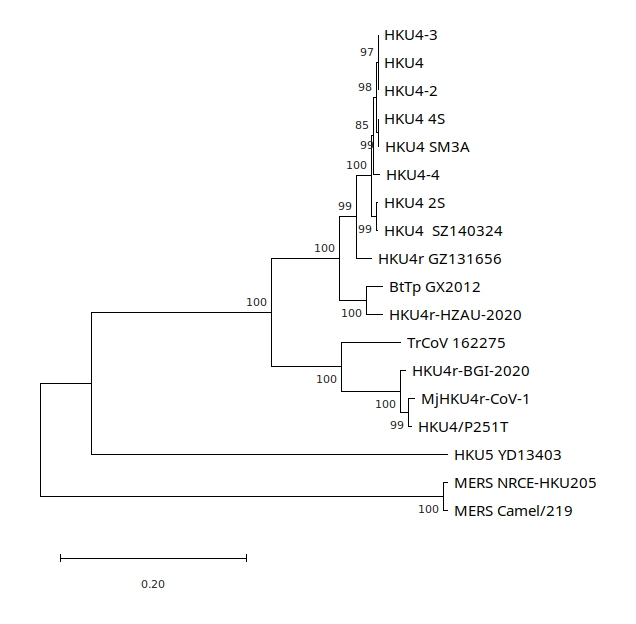

Supp. Fig. 2. Maximum likelihood tree of spike gene. Tree drawn to scale, with branch lengths measured in the number of substitutions per site, bootstrap values >70% are shown. Tree rooted on midpoint and displayed using MEGA11. Accession numbers and names in Supp. Info. 3.

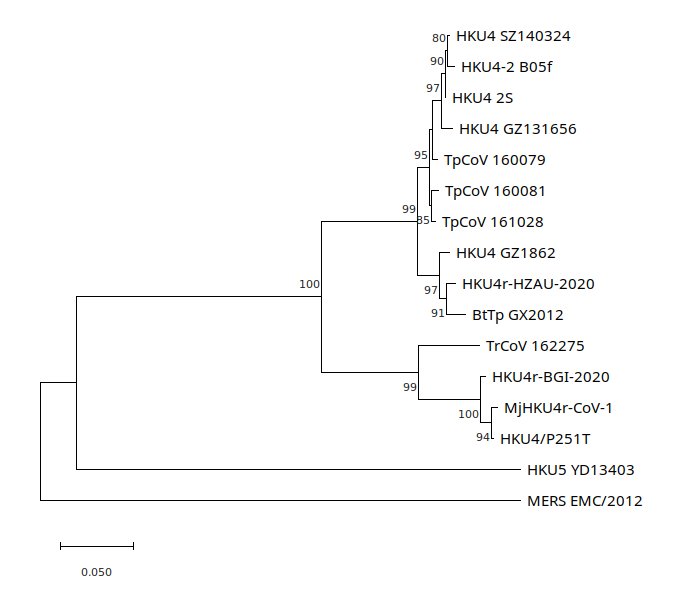

Supp. Fig. 3. Maximum likelihood tree of spike protein sequences of 14 HKU4-related CoVs, HKU5, and MERS. Tree drawn to scale, with branch lengths measured in the number of substitutions per site, bootstrap values >70% are shown. Tree rooted on midpoint and displayed using MEGA11. Accession numbers and names in Supp. Info. 3.

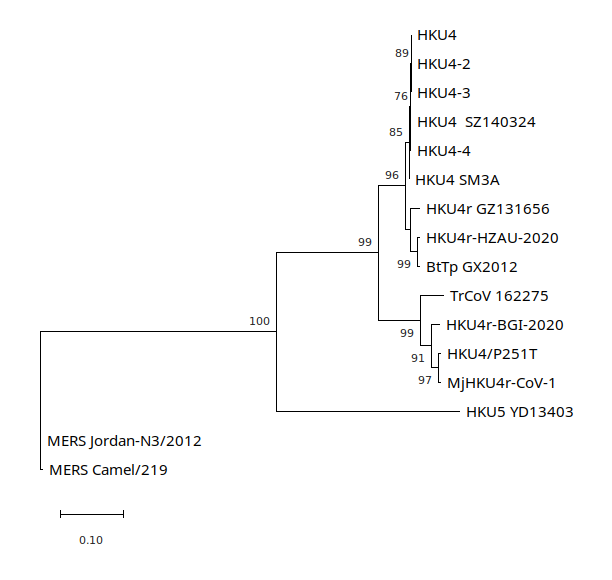

Supp. Fig. 4. HKU4-related CoV N gene maximum likelihood tree. Tree drawn to scale, with branch lengths measured in the number of substitutions per site, bootstrap values >70% are shown. Tree rooted on midpoint and displayed using MEGA11. Accession numbers and names in Supp. Info. 3.

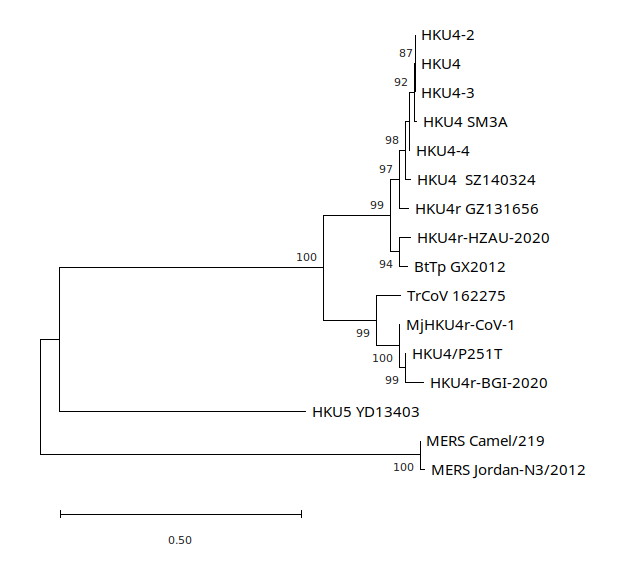

Supp. Fig. 5. HKU4-related CoV ORF3-5 gene region maximum likelihood tree. Tree drawn to scale, with branch lengths measured in the number of substitutions per site, bootstrap values >70% are shown. Tree rooted on midpoint and displayed using MEGA11. Accession numbers and names in Supp. Info. 3.

#### Spike Protein

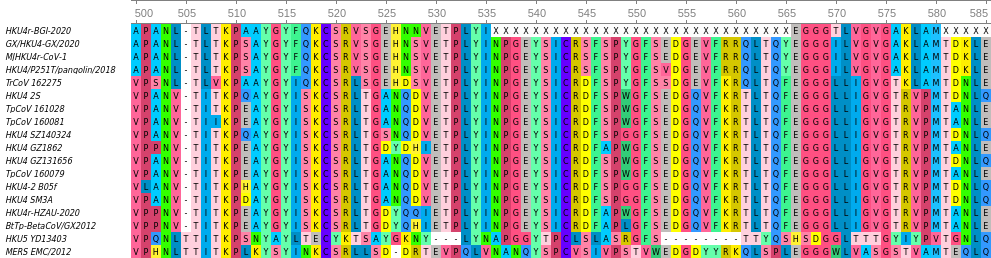

Supp. Fig. 6. Receptor binding motif section of spike protein after alignment of spike protein sequences with the highest homology to HKU4r-BGI-2020 (consensus reference numbering).

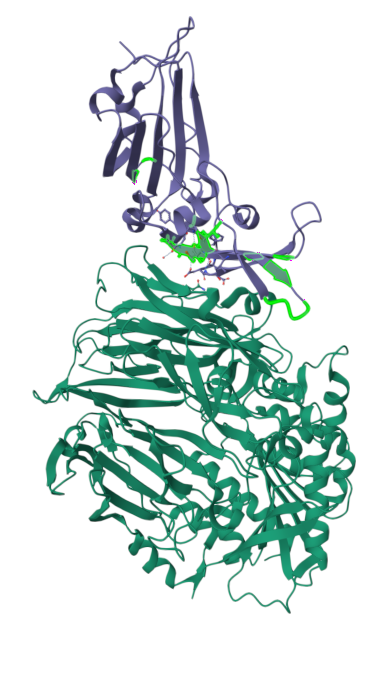

Supp. Fig. 7. 3D structure of the S1 domain of S protein of BtCoV HKU4 (purple) in complex with hDPP4 (dark green). Seventeen amino acids in the receptor binding motif of HKU4 with differences to HKU4r-BGI-2020 are highlighted in bright green.

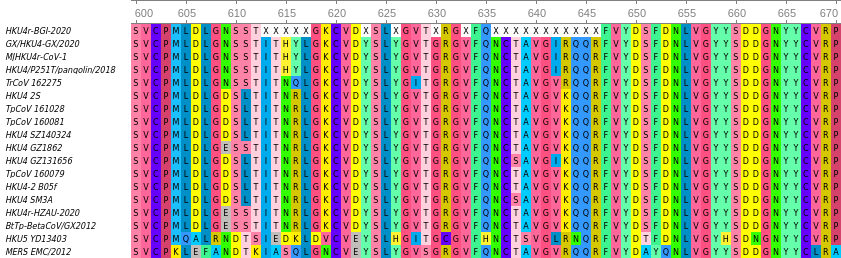

Supp. Fig. 8. Amino acid positions 600-670 (consensus reference numbering) for alignment of spike protein sequences to HKU4r-BGI-2020. Includes coverage of RQQR putative furin cleavage motif (Chen et al., 2023) at positions 643-646.

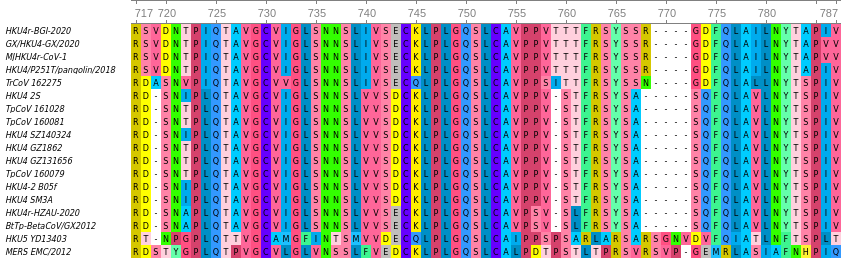

Supp. Fig. 9. Alignment of spike protein sequences to HKU4r-BGI-2020 displaying amino acid positions 717-787 (consensus reference numbering) including MERS (RSVR) and HKU5 (RSAR) S1/S2 junction furin cleavage motifs at positions 765-768.

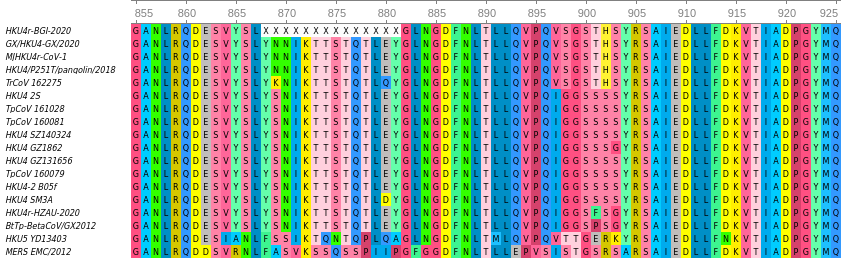

Supp. Fig. 10. Alignment of spike protein sequences to HKU4r-BGI-2020 displaying amino acid positions 855-925 (consensus reference numbering) including S2’ MERS (RSAR) furin cleavage motif at positions 902-905.

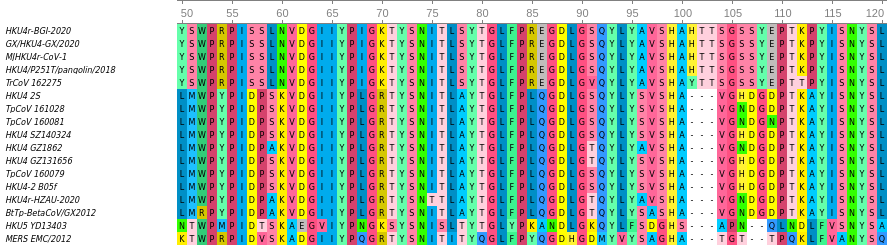

Supp. Fig. 11. Alignment of spike protein sequences to HKU4r-BGI-2020 displaying amino acid positions 50-120 (consensus reference numbering). HKU4r-BGI-2020, PCoV HKU4-P251T, PCoV MjHKU4r-CoV-1 and *Tylonycteris robustula* bat CoV 162275 contain a three amino acid insert in the NTD of the S protein not found in other known HKU4-related CoV sequences, HKU5-CoV or MERS-CoV.

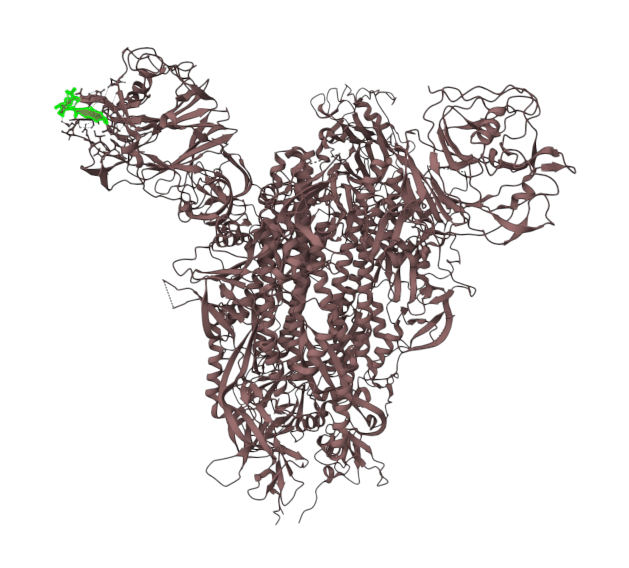

Supp. Fig. 12. 3D structure of the MERS-CoV S protein PDB: 7X27 viewed using Mol 3D viewer. Sequence GHA---TGT--T (AA positions 90-95, dashes indicating location of gaps relative to HKU4r-BGI-2020) in NTD of the S1 domain of the S protein highlighted in green. The three amino acid insert HTT in HKU4r-BGI-2020, PCoV HKU4-P251T, PCoV MjHKU4r-CoV-1 and YTT in TrCoV 162275 S protein sequences relative to other HKU4-related CoVs occurs in an external position of the structure.

#### Selection Pressure

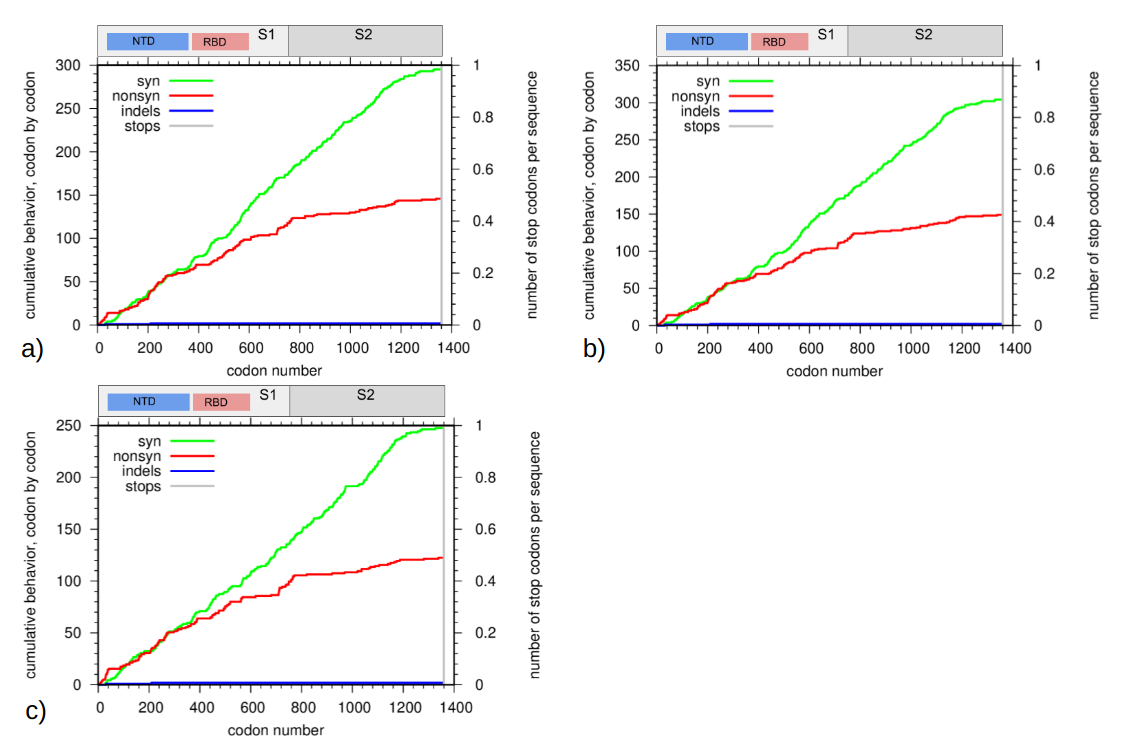

Supp. Fig. 13. Cumulative dN/dS plots showing indels and stop codons for spike protein sequences for a) PCoV HKU4-P251T and TrCoV 162275; b) MjHKU4r-CoV-1 and TrCoV 162275; and c) HKU4-BGI-2020 and TrCoV 162275, plotted using SNAP (Korber, 2002). Missing amino acid sections in HKU4-BGI-2020 are located at positions 185-203; 404-420; 528-557; 574-583; 628-638; 856-869 and 976-1011.

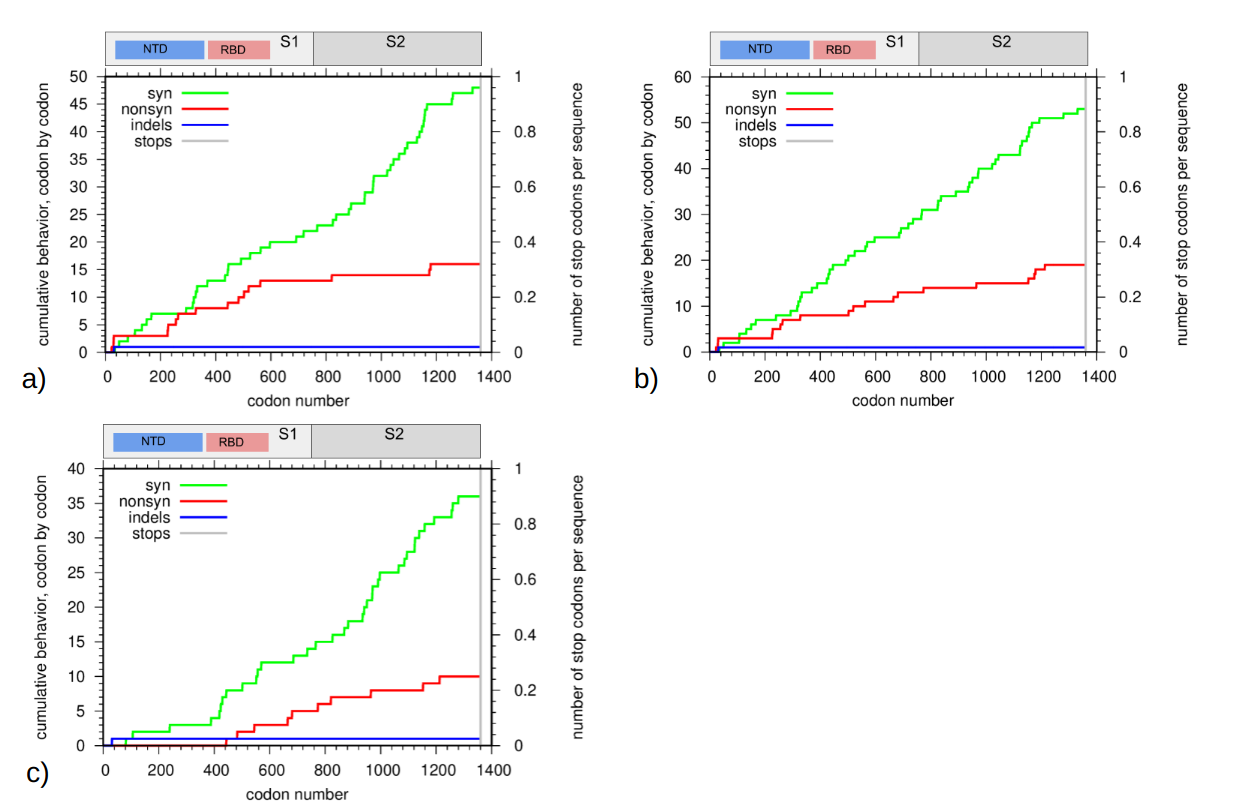

Supp. Fig. 14. Cumulative dN/dS plots showing indels and stop codons for spike protein sequences from a) PCoV HKU4-P251T and HKU4r-BGI-2020; b) MjHKU4r-CoV-1 and HKU4r-BGI-2020; and c) PCoV HKU4-P251T and MjHKU4r-CoV-1. Plotted using SNAP (Korber, 2002). Missing amino acid sections in HKU4-BGI-2020 are located at positions 185-203; 404-420; 528-557; 574-583; 628-638; 856-869 and 976-1011.

#### Host analysis

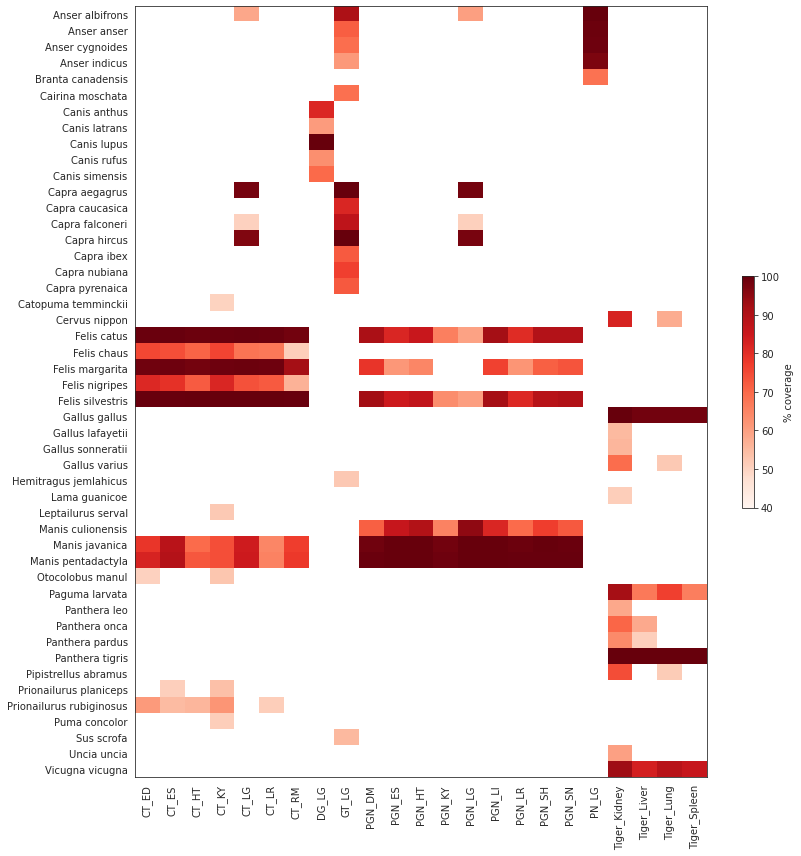

Supp. Fig. 15. Mitochondrial alignments of selected datasets in BioProject PRJNA747757 to all mitochondrial genomes on NCBI using minimap2. A 50% genome coverage cutoff was applied prior to plotting.

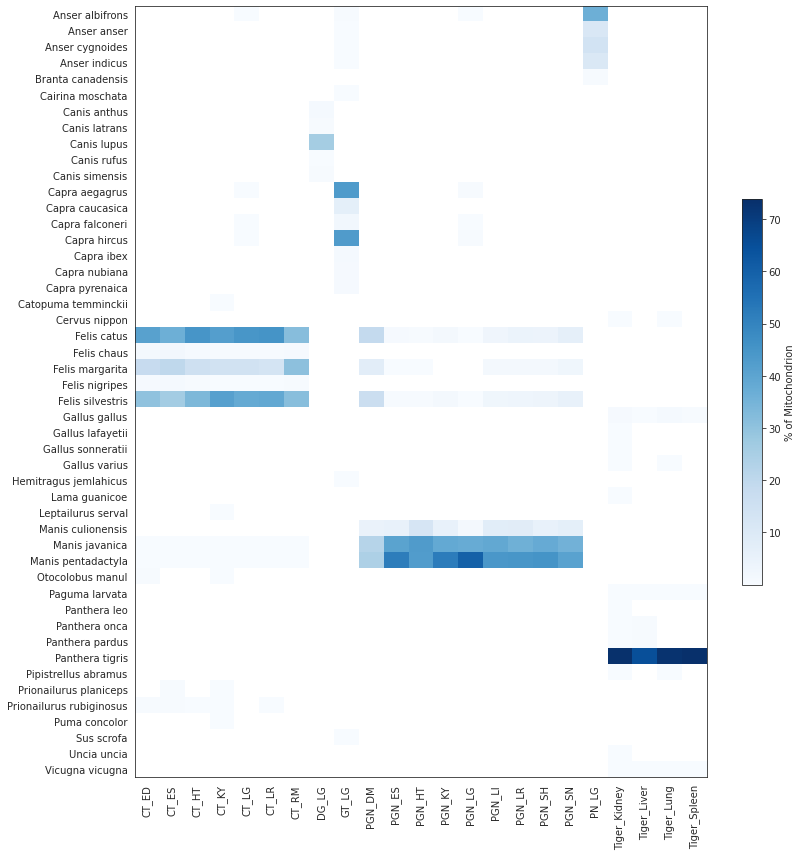

Supp. Fig. 16. Percent of mitochondrial alignments of selected datasets in BioProject PRJNA747757 to all mitochondrial genomes on NCBI using minimap2 with a greater than 10% coverage. A 50% genome coverage cutoff was applied prior to plotting.

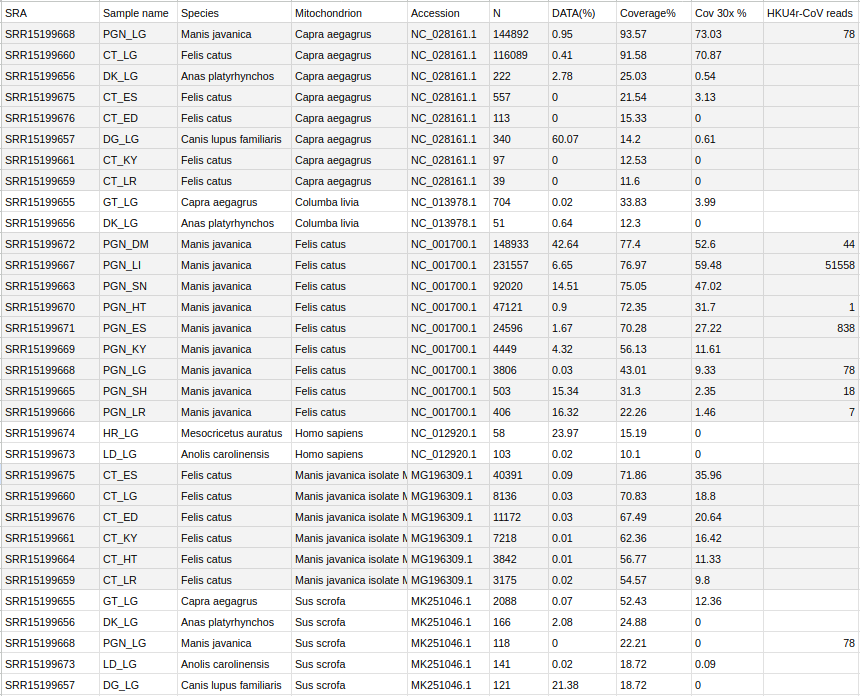

Supp. Table 1. Contaminating mitochondrial genomic sequences in selected NGS datasets in BioProject PRJNA747757, with DATA(%) indicating percentage of all mitochondrial alignments per SRA sample.

#### Contaminating Viruses

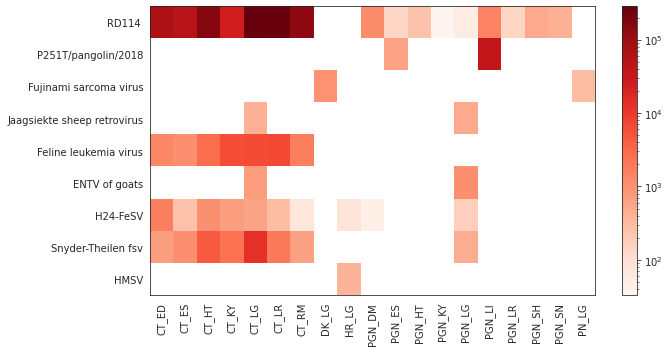

Supp. Fig. 17. Read count for reads matching viruses identified in BioProject PRJNA747757, using a virus minimum coverage of 10%. Displayed using a log scale.

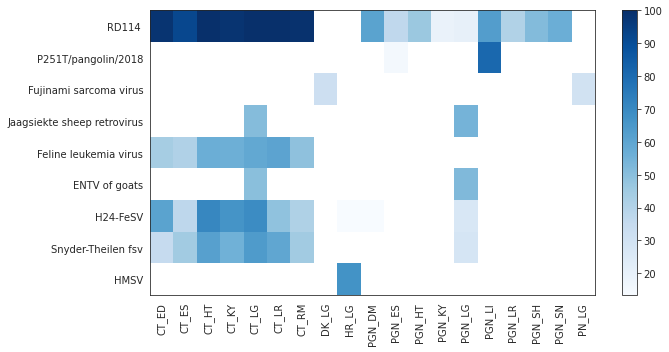

Supp. Fig. 18. Per sample coverage percentage of viruses identified in BioProject PRJNA747757. A virus minimum coverage of 10% was used.

#### Host analysis of “clade b” HKU4-related CoVs

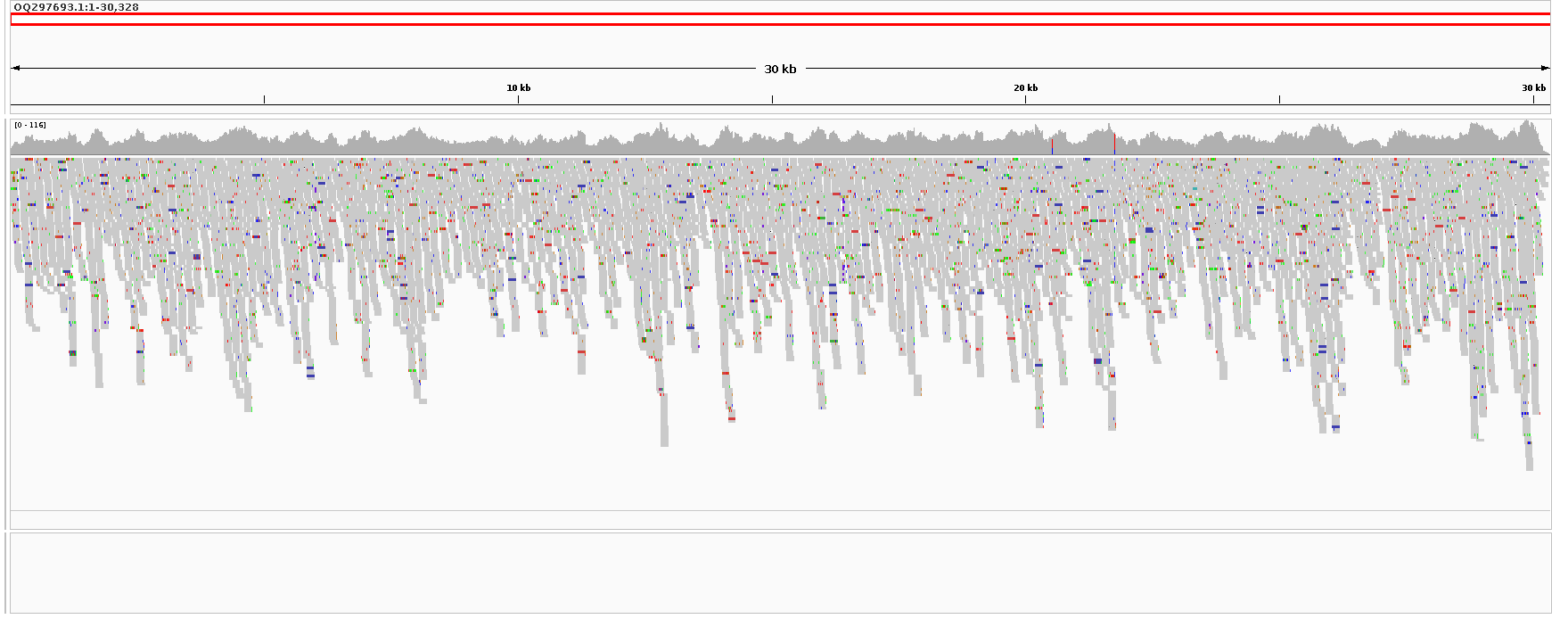

Supp. Fig. 19. Alignment of reads is NGS dataset for sample HKU4-GX (SRR22936420) in BioProject PRJNA901878 to MAG: Pangolin coronavirus HKU4 isolate GX/HKU4-GX/2020 (OQ297693.1). Aligned using minimap2, displayed using IGV.

| Accession | N | Avg depth | Median | Coverage% | Cov 4x % | Cov 10x % | Cov 30x % | Cov 100x % |
| --- | --- | --- | --- | --- | --- | --- | --- | --- |
| OQ297693.1 | 14404 | 71.06 | 68 | 100 | 99.98 | 99.84 | 99.07 | 11.25 |

Supp. Table 2. Read mapping statistics for alignment of HKU4-GX (SRR22936420) in BioProject PRJNA901878 to MAG: Pangolin coronavirus HKU4 isolate GX/HKU4-GX/2020.

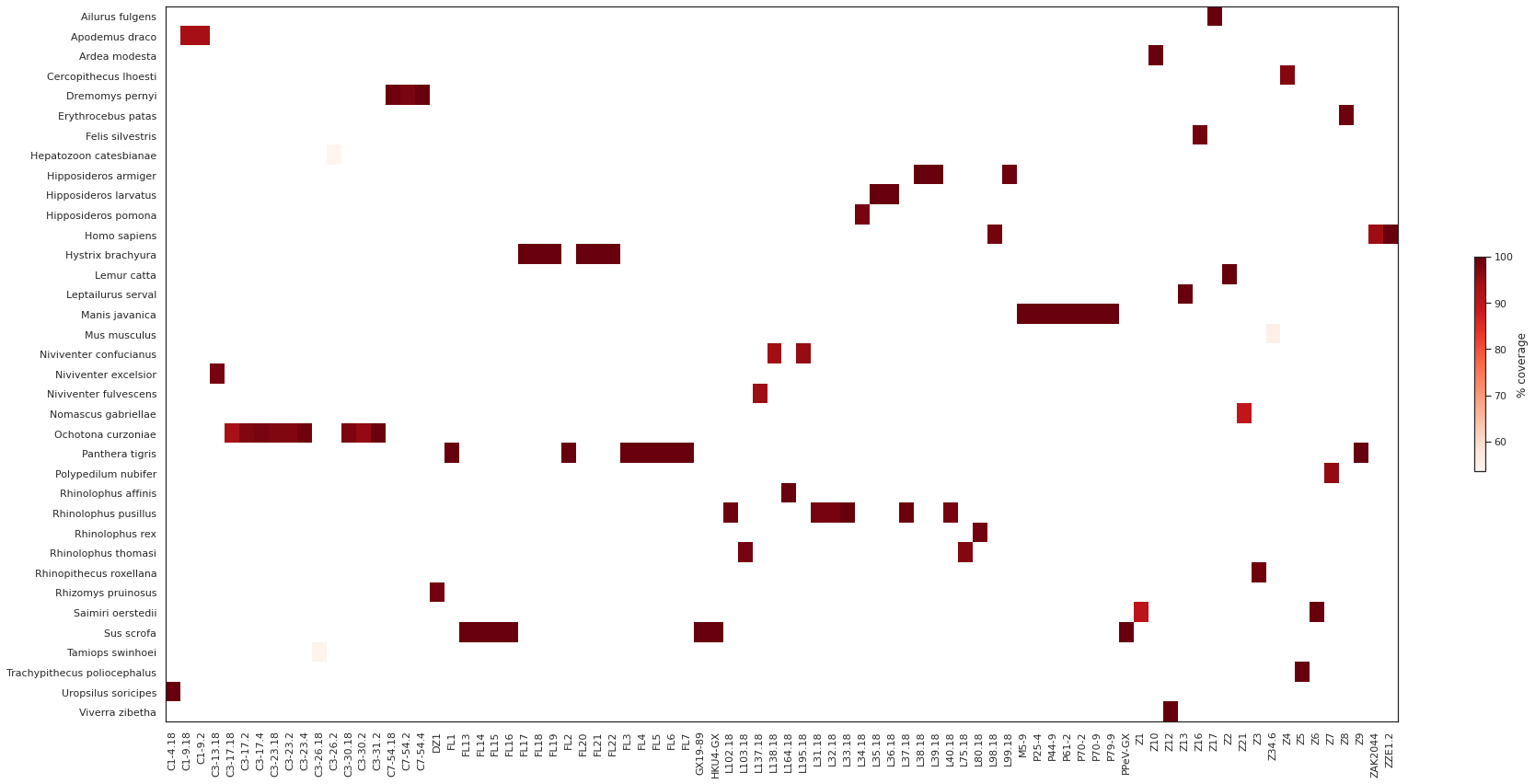

Supp. Fig. 20. Mitochondrial alignments of 86 datasets in BioProject PRJNA901878 to all mitochondrial genomes on NCBI using minimap2. Only datasets with 50% genome coverage or greater are plotted.

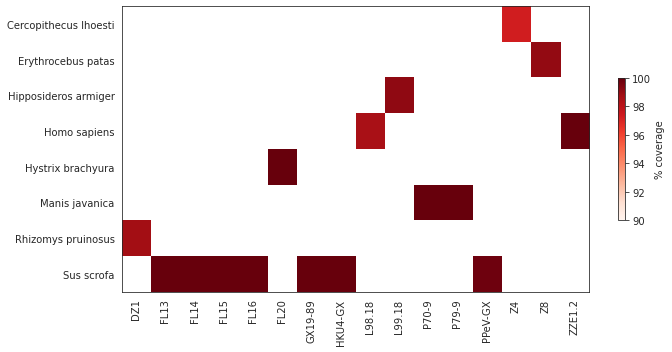

Supp. Fig. 21. Mitochondrial alignments of selected datasets in BioProject PRJNA901878 to all mitochondrial genomes on NCBI using minimap2. A 50% genome coverage cutoff was applied prior to plotting.

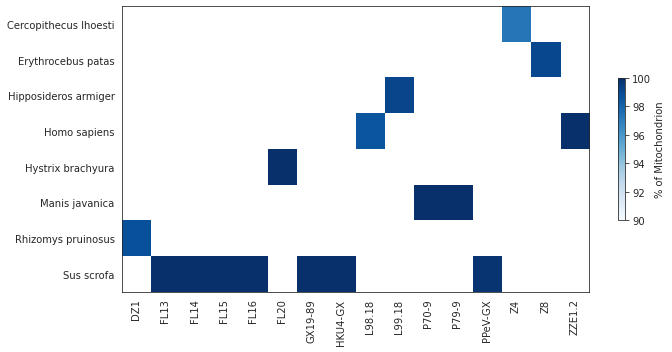

Supp. Fig. 22. Percentage of total mitochondrial alignments of selected datasets in BioProject PRJNA901878 to all mitochondrial genomes on NCBI using minimap2. A 50% genome coverage cutoff was applied prior to plotting.

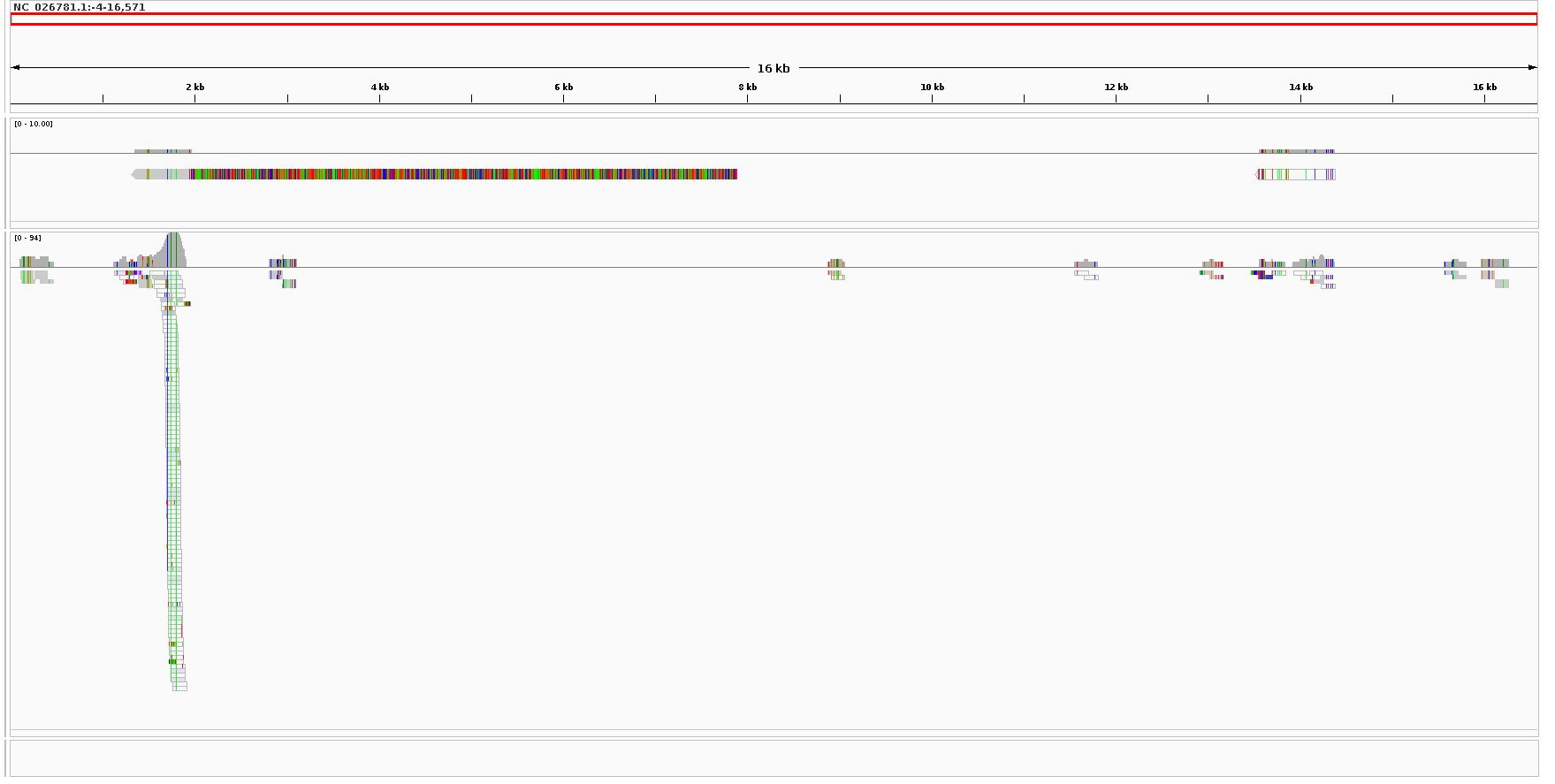

Supp. Fig. 23. *De novo* assembled contigs (top track) and reads in dataset HKU4-GX (SRR22936420) in BioProject PRJNA901878 aligning to the *Manis javanica* isolate T298 mitochondrial genome (NC_026781.1).

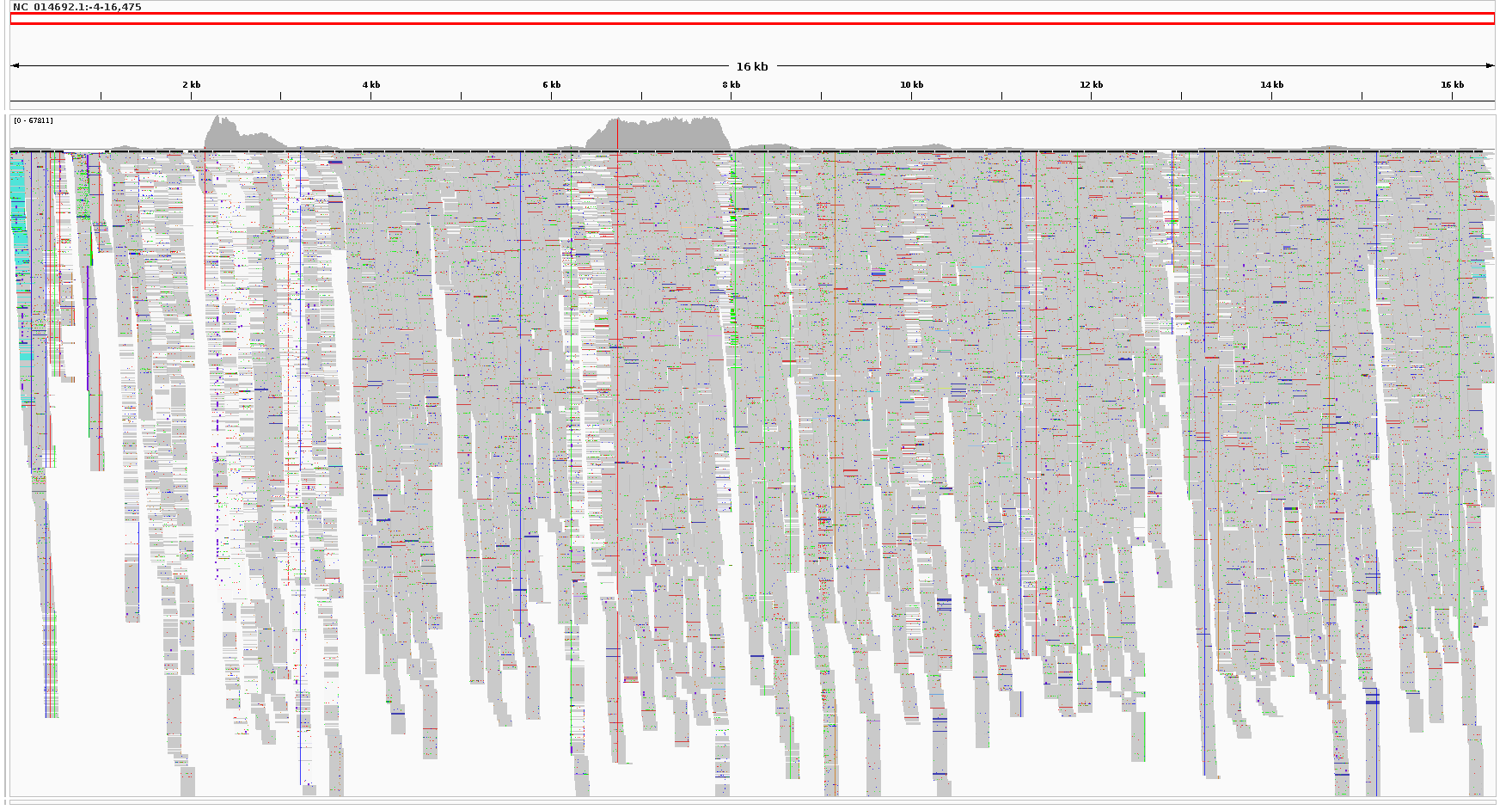

Supp. Fig. 24. Reads in NGS dataset for sample HKU4-GX (SRR22936420) in BioProject PRJNA901878 aligning to the *Sus scrofa taiwanensis* mitochondrial genome (NC_014692.1). Top track read depth scale is 0-67811. Average read depth is 515.49 with 100% genome coverage. Displayed in IGV with options to downsample reads and remove duplicates.

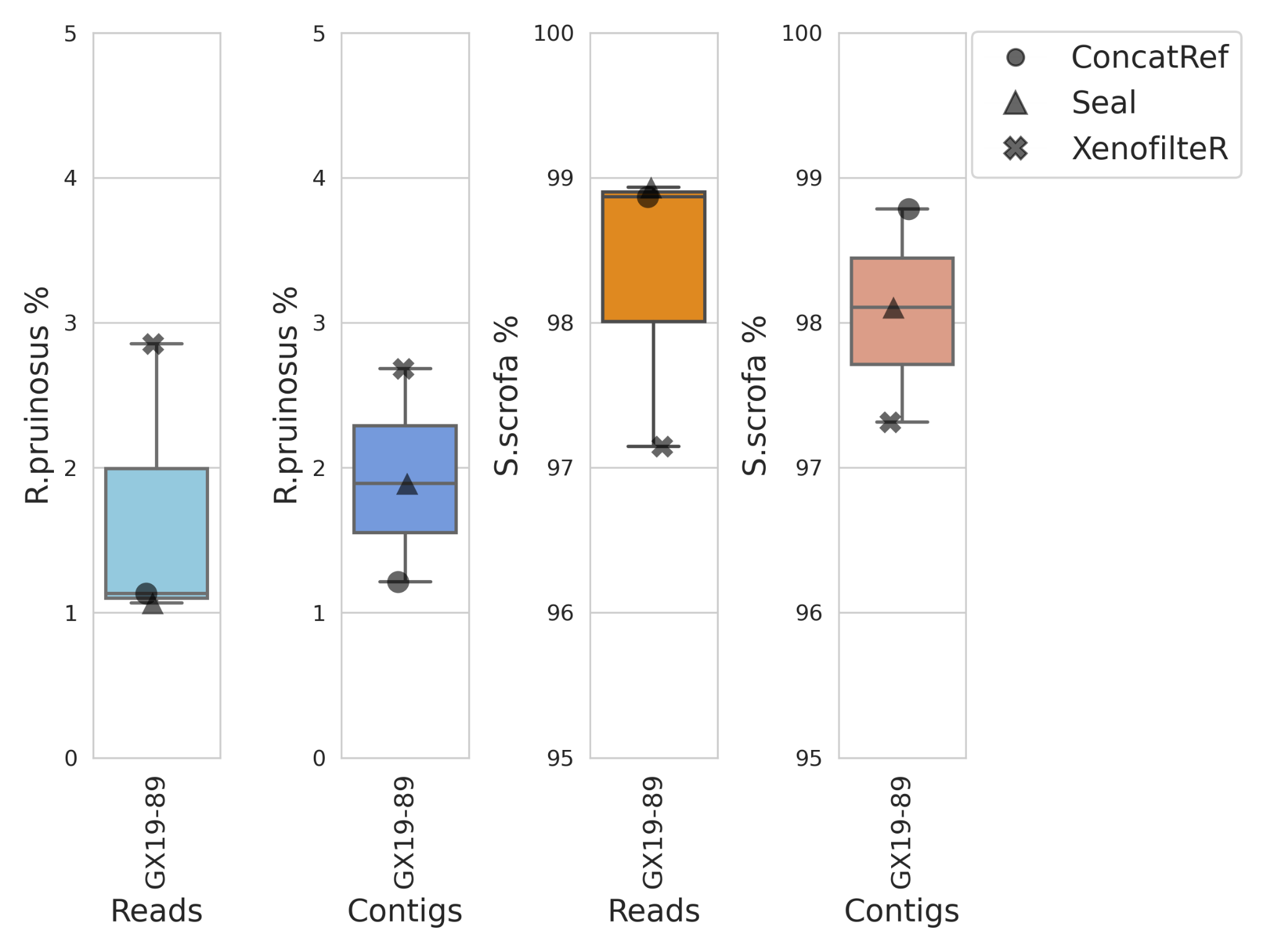

Supp. Fig. 25. *Rhizomys pruinosus* and *Sus scrofa* genomic percentage content for reads and *de novo* assembled contigs in RNA-Seq dataset SRR22936497 from sample GX19-89 in BioProject PRJNA901878 using three independent methods.

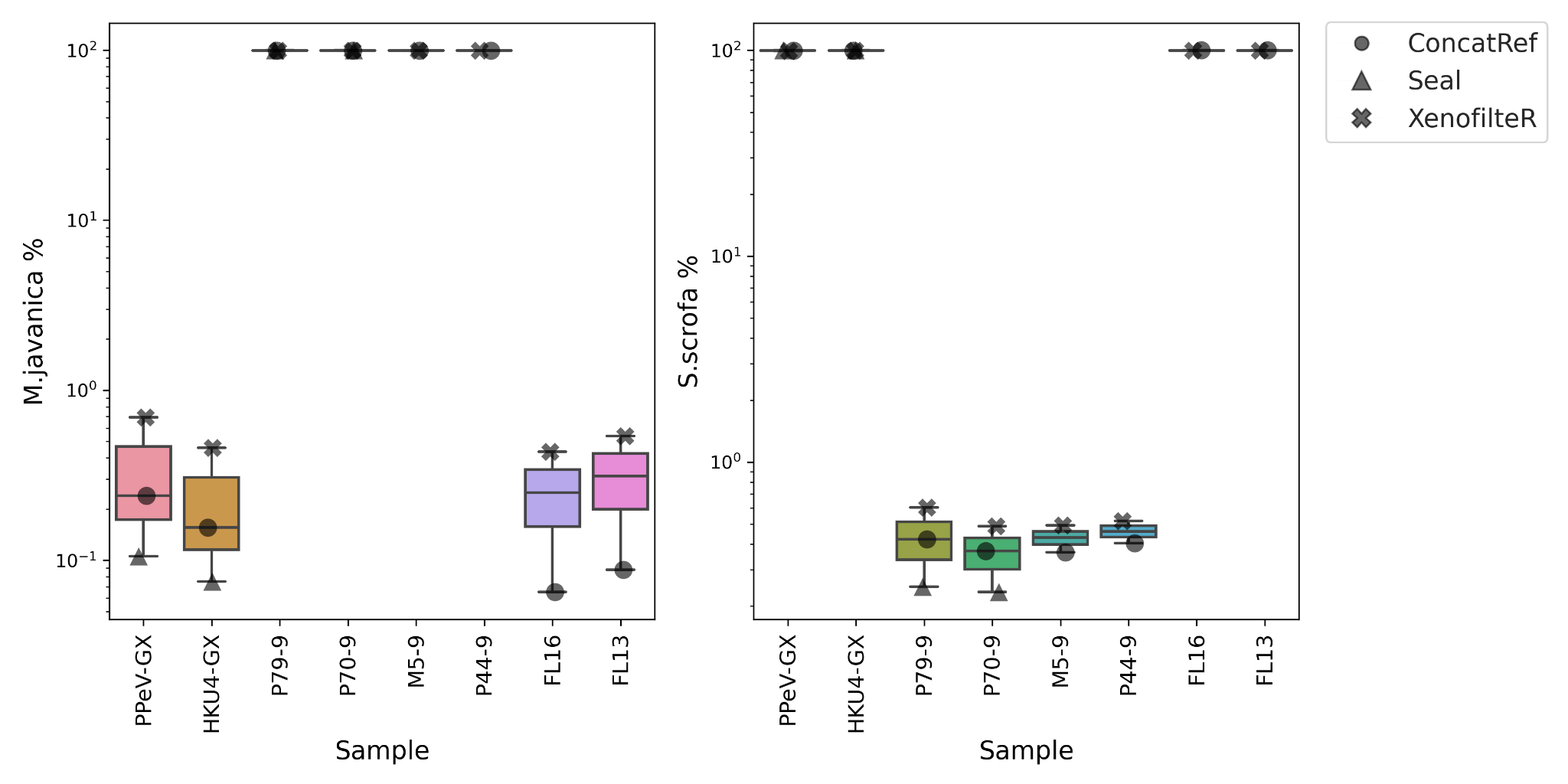

Supp. Fig. 26. *De novo* assembled contigs for selected datasets in BioProject PRJNA901878 were filtered to include only those 300-nt or longer then aligned to *Manis javanica* and *Sus scrofa* genomes using three methods. Percentage of contigs best aligning to *M. javanica* or *S. scrofa* are shown in log scale.

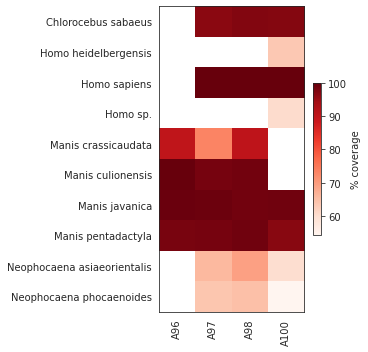

Supp. Fig. 27. Alignment of four SRA datasets in BioProject PRJCA002517 containing HKU4-related CoVs to all mitochondrial genomes on NCBI. Only genomes with greater than 50 percent coverage are shown.

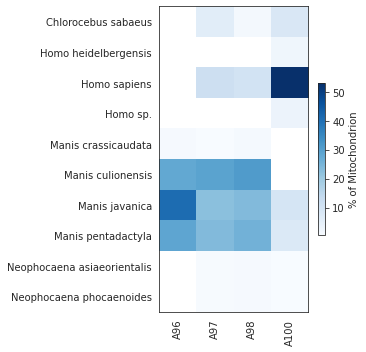

Supp. Fig. 28. Alignment of four CRA datasets in PRJCA002517 containing HKU4-related CoVs to all mitochondrial genomes on NCBI. Read counts were divided by total read counts matching mitochondrial genomes where coverage was greater than 10% to give percentages shown. Only genomes with greater than 50 percent coverage are plotted.

| Run | Sample | Isolate | N | Avg depth | Median | Cov 1x% | Cov 4x % | Cov 10x % | Cov 30x % | Cov 100x % |
| --- | --- | --- | --- | --- | --- | --- | --- | --- | --- | --- |
| CRR477157 | A100 | MjHKU4r-CoV-4 | 64506 | 314.51 | 242 | 99.99 | 99.96 | 99.96 | 99.43 | 88.62 |
| CRR477156 | A98 | MjHKU4r-CoV-3 | 701 | 3.41 | 3 | 83.89 | 37.54 | 5.49 | 0 | 0 |
| CRR477155 | A97 | MjHKU4r-CoV-2 | 10062 | 49.03 | 37 | 99.96 | 99.22 | 94.61 | 62.09 | 9.64 |
| CRR477154 | A96 | MjHKU4r-CoV-1 | 2538 | 12.18 | 11 | 99.13 | 92.56 | 57.05 | 2.38 | 0 |

Supp. Table 3 Summary statistics for read alignments to MjHKU4r-CoV-1 to 4 for four datasets in BioProject PRJCA002517.

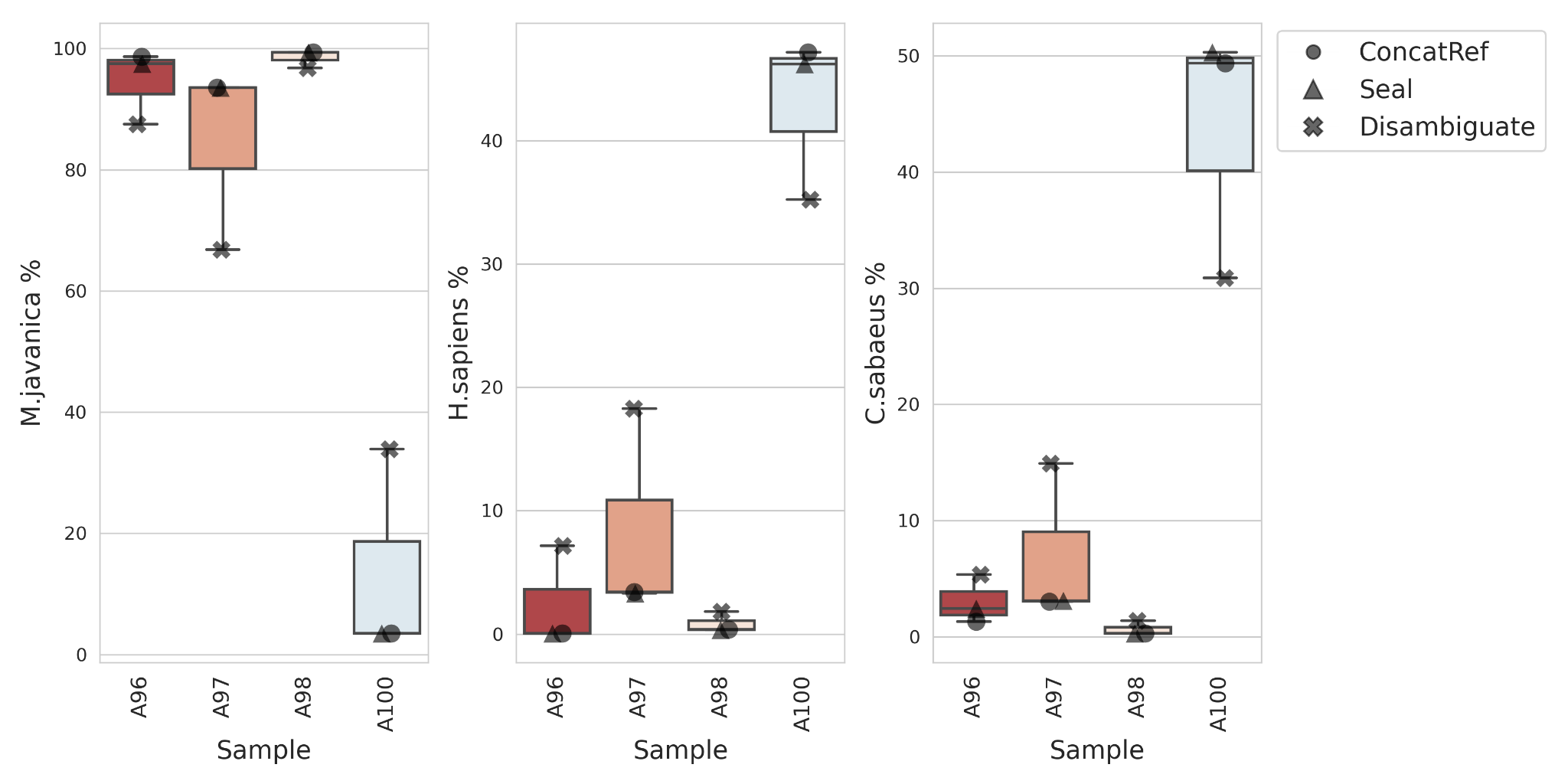

Supp. Fig. 29. PRJCA002517 dataset full genome read alignments. Reads in four datasets documented as generated from pangolin samples (Shi et al., 2023) were aligned to full *Manis javanica*, *Homo sapiens*, and *Chlorocebus sabaeus* genomes for using ConcatRef (Jo et al., 2017), Seal (Bushnell, 2022), and Disambiguate (Valentine, 2020) workflows. Percentages of reads mapping to each genome were plotted separately.

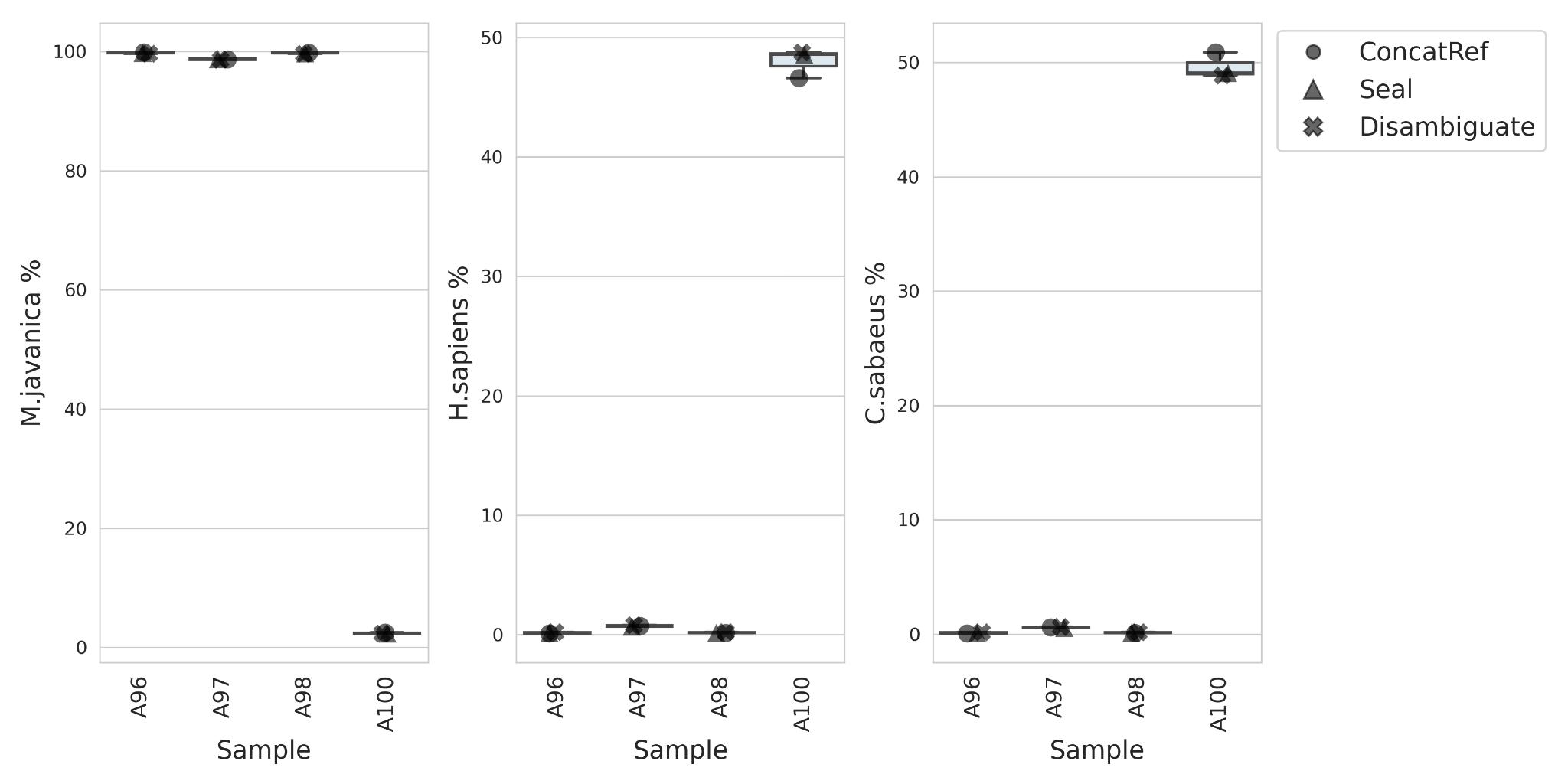

Supp. Fig. 30. PRJCA002517 dataset full genome contig alignments. *De novo* assembled contigs in four datasets documented as generated from pangolin samples (Shi et al., 2023) were aligned to full *Manis javanica*, *Homo sapiens*, and *Chlorocebus sabaeus* genomes for using ConcatRef (Jo et al., 2017), Seal (Bushnell, 2022), and Disambiguate (Valentine, 2020) workflows. Percentages of reads mapping to each genome were plotted separately.

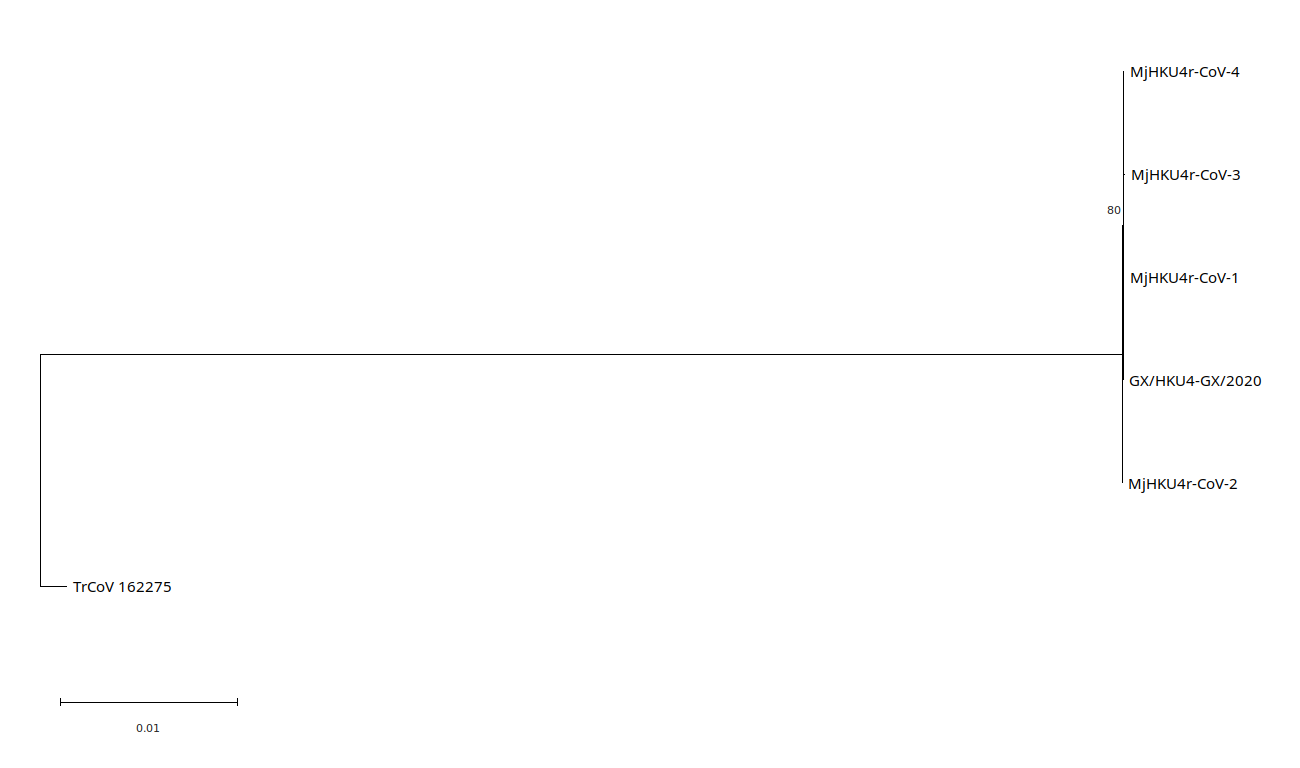

Supp. Fig. 31. Maximum likelihood tree for full genomes for MjHKU4r-CoV-1 and variants and *Tylonycteris robustula* CoV 162275. Generated using MEGA11 using a GTR+I model with 100 bootstrap replicates. Tree drawn to scale, with branch lengths measured in the number of substitutions per site. Tree rooted on TrCoV 162275 and displayed using MEGA11. Bootstrap values >70% are shown, accession numbers and names in Supp. Info. 3.
